# Profibrotic priming of airway cell types and drug responses in early-stage idiopathic pulmonary fibrosis

**DOI:** 10.1101/2022.03.09.483638

**Authors:** Robert Lorenz Chua, Carmen Veith, Marc A. Schneider, Katharina Jechow, Elizabeth Chang Xu, Michael Kreuter, Agnes W. Boots, Roland Eils, Nicolas C. Kahn, Christian Conrad

## Abstract

Early genetic studies hinted the role of airway epithelial cells in the development of idiopathic pulmonary fibrosis (IPF), while recent single-cell RNA sequencing (scRNA-seq) atlases utilized explant IPF lungs and therefore represent late-stage disease. Here, we used air liquid interface (ALI) cultures of primary cells taken from the subsegmental bronchi of newly diagnosed IPF patients, reflecting early-stage fibrosis, to interrogate the transcriptional landscape of the airway mucosa. Profiling of 129,986 cells identified a shared proinflammatory state in epithelial cells and an early activation state of fibroblasts. Moreover, IPF basal cells initiated awry repair mechanisms and primed the airway mucosa for TGF-β activation. Treatment with nintedanib, pirfenidone, both established antifibrotic drugs, and saracatinib, an Src kinase inhibitor that can limit IPF progression, only significantly affected certain IPF signatures. This study provides insight into the early disease mechanisms of IPF and may serve as a resource to further investigate pharmacological inhibition effects.

## Introduction

Idiopathic pulmonary fibrosis is a rare, chronic interstitial lung characterized by progressive parenchymal scarring, extracellular matrix (ECM) deposition, progressively impaired lung function, and high mortality. It is associated with a poor prognosis with a median survival of 3-5 years after diagnosis with an incidence of about 0.09-1.30 cases per 100,000 individuals (*1*–*3*). This disease is defined by the presence of radiological and/or histopathological pattern of usual interstitial pneumonia without any attributable etiology; this is reflected by the extensive remodeling of the lung architecture through the presence of fibrotic foci, honeycomb cysts, dilated bronchi, and thickened alveolar septa. The development of IPF manifest at the distal ends of the lung and progressively develops upwards through the entire lung tissue (*1*).

Previous studies using bulk tissue sequencing technologies revealed genetic mutations (e.g. *MUC5B, TERT*) (*4, 5*) and altered transcriptomes (e.g. *COL1A1, MMP7, SPP1, KRT5, MUC5B, DNAI1*) (*6, 7*) that define IPF as a disease. These highlight the significance and alterations of the lung epithelium and mesenchyme in the development of IPF. However, a major caveat of bulk sequencing is the loss of cellular resolution, and pathological mechanisms cannot be directly linked to specific cell populations. Early studies applying single-cell RNA sequencing (scRNA-seq) on explant IPF lungs have low numbers of epithelial cells or focused on macrophages (*8, 9*). Later IPF scRNA-seq atlases provided a broader understanding of different cellular populations (e.g. macrophages, fibroblasts, endothelial, and epithelial cells) and their pathogenic roles in fibrosis (*10*–*12*).

Nevertheless, while early studies hinted that airway epithelial cells may play an important role in the development of IPF (*5*–*7*), more effort into this warranted. Moreover, previous studies analyzed the end stages of IPF, hindering the identification of potential “early-stage” cellular phenotypes that could be targeted by drugs. To address these limitations, we performed scRNA-seq on air-liquid interface (ALI) cultured primary human bronchial epithelial cells (HBEC) that were taken from the patients with newly diagnosed and untreated IPF via bronchoscopic microsampling, thus, representing early-stage IPF. ALI-cultured cells were treated with nintedanib and pirfenidone, antifibrotic compounds currently recommended to treat IPF (*13*). Additionally, the use of a Src kinase inhibitor, saracatinib, was also investigated as these kinases are implicated in cellular events involved in IPF like cellular proliferation, myofibroblast differentiation, and epithelial-to-mesenchymal transition (EMT) (*14*–*16*). While the effects of these drugs have been studied in fibroblasts and myofibroblasts, their effects on epithelial cells are still unknown.

In this study, we identified broad changes in the cellular composition of cultured airway mucosa cells, with an expansion of fibroblasts and reduction of basal cells in IPF using scRNA-seq. We also identified specific pathways upregulated in IPF cell populations and aberrant transcriptomic signatures in epithelial cells and fibroblasts that we further validated on lung biopsies from newly diagnosed IPF patients. Furthermore, these early-stage pathogenic signatures were not fully rescued by treatment with nintedanib, pirfenidone nor saracatinib.

## Results

### Cellular diversity of the early-stage IPF airway epithelium

To investigate differences in cellular composition and transcriptomic phenotypes of the medial conducting airways between non-fibrotic control and early-stage IPF, we performed 3’ scRNA-seq on primary ALI-cultured HBECs. A minimally invasive procedure was used to obtain HBECs in the endobronchial lining fluid (ELF) from the subsegmental branches of the lung from both patient groups. These primary cells were differentiated in an ALI-culture before being collected for scRNA-seq. Mature ALI-cultures were also treated with nintedanib, pirfenidone, or saracatinib to assess their effects on IPF cells (Fig. 1A).

**Figure 1.**
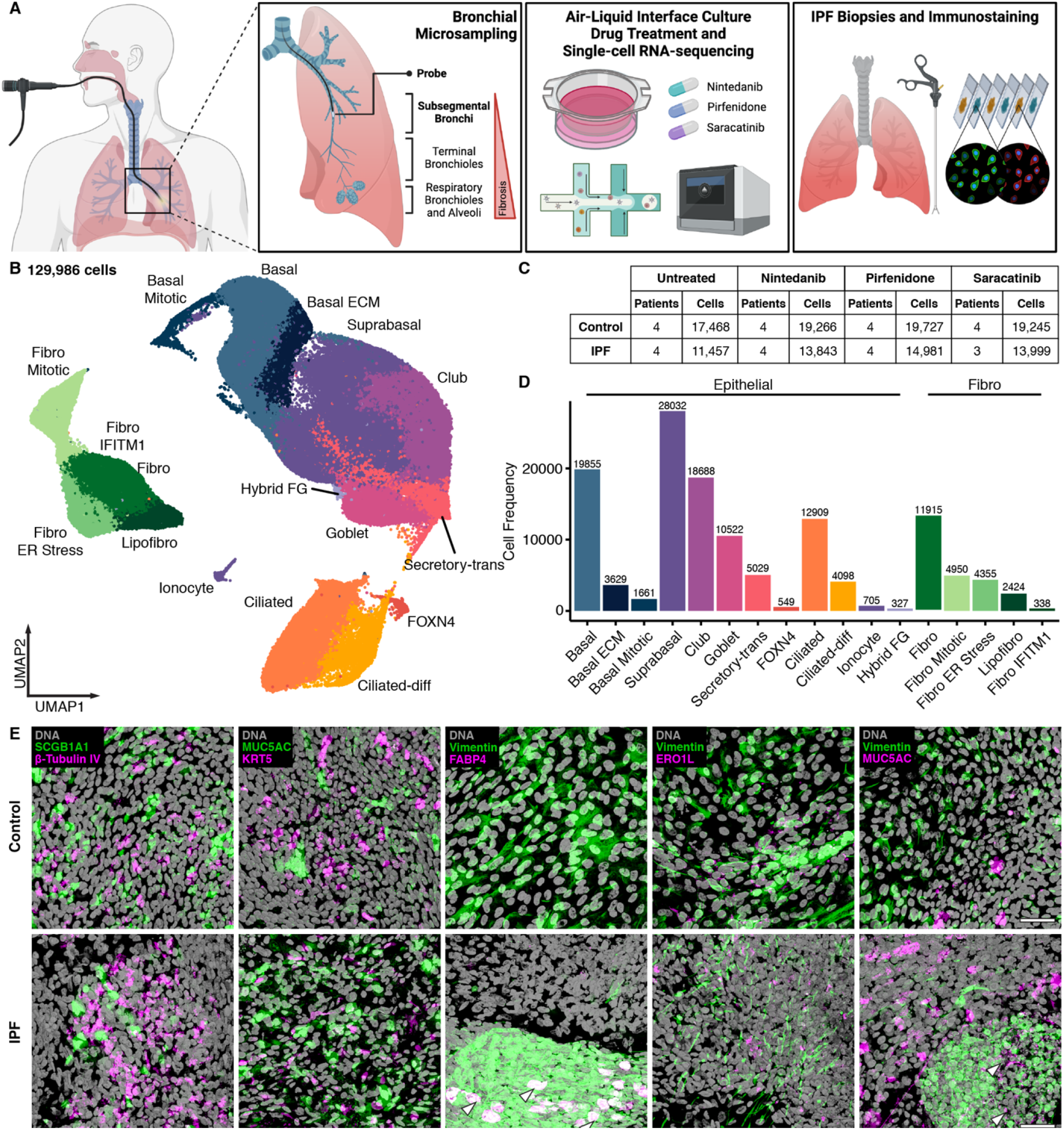
Experimental and dataset overview of the study. (**A)** Epithelial lining fluid (ELF) was obtained by bronchial microsampling from control and newly diagnosed IPF patients. HBECs were grown in an ALI-culture and treated with nintedanib, pirfenidone or saracatinib. Cells were harvested and captured for scRNA-seq and immunofluorescence. (**B)** UMAP embedding of 129,986 scRNA-seq transcriptomes from 17 different airway cell types/states from all patients and treatments. (**C)** Patient cohort overview showing the number of patients and cells sequenced per disease and treatment categories. (**D)** Overall cellular frequencies of each cell type and state. (**E)** Immunofluorescent images stained for markers of epithelial and mesenchymal cell types in ALI-cultures of control and IPF patients. FABP4^+^ fibroblasts are highlighted with white arrowheads. Fibroblast clusters intermingled with MUC5AC are highlighted with white arrowheads. Scale bar = 50μm. Validation cultures: control (n = 2); newly diagnosed IPF (n = 2). Abbreviations: Fibro – Fibroblast, Lipofibro – Lipofibroblast, FG – Fibroblast/Goblet.

In total, we generated a collection of 129,986 transcriptomes from 8 patients and a total of 31 samples (Fig. 1B, C). We identified 17 cell types and states from the entire dataset depicting previously described epithelial cells from the airway mucosa (basal, suprabasal, club, goblet, ciliated, ionocytes, and FOXN4 cells) (*17, 18*) that accounted for more than half of the entire cellular population, while the rest consisted of a large number of fibroblast-like cells (Fig. 1B, D; fig. S1). Immunofluorescent staining of ALI-cultures from control and IPF patients for different cell type markers validate the presence of the major cell types in our model system (Fig. 1E, fig. S2). Comparisons with previous IPF scRNA-seq datasets additionally confirmed the high concordance of our system with primary tissue (*10*–*12*) (fig. S3A-C).

We identified subclusters of different cell types through unsupervised clustering and characterized their differences on the transcriptomic level with different marker genes, gene ontology (GO) term enrichment, and pseudo-temporal dynamics (Fig. 1B, D; fig. S1, 3D-F). Basal cells were split into three clusters: basal, basal ECM, and basal mitotic. Basal cells were in a state of protein translation and biogenesis (fig. S1). On the other hand, basal ECM cells highly expressed structural genes like *FN1* and *MMP9* and were also enriched for GO terms like ECM organization and wound healing (fig. S1). Basal mitotic expressed mitosis genes such as *CDK1* and *TOP2A* (fig. S1). Ciliated cells were split into two states and defined by their maturation status as shown by differences in expression of cilium genes, GO terms for differentiation and development, and cellular trajectories (fig. S1, 3D-F). We also observed two transitionary states: suprabasal and secretory-trans cells (Fig. 1B, D; fig. S1, 3D-F). Suprabasal cells are early progenitors derived from basal cells that later differentiate into airway epithelial cells (*19, 20*). They express serpin family genes, like *SERPINB3* and *SERPINB13*, and *KRT19* (*20*) (fig. S1). On the other hand, secretory-trans cells represent secretory cell types transitioning into ciliated cells (*19*). This is shown by the reduced expression of secretory markers and their proximity to the ciliated cell clusters on the manifold (fig. S1, 3D-F). Finally, we also observed a small population of epithelial cells that are mostly found in IPF samples with a mixed gene signature of goblet cells and fibroblasts that we called “hybrid FG” (fig. S3G, H). In IPF ALI-cultures, we detected fibroblast foci co-expressing vimentin and MUC5AC (Fig. 1E; fig. S3H).

While ALI-cultures promote the growth and differentiation of airway epithelial cells, we observed a large population of fibroblast-like cells in our dataset. All five populations expressed fibroblast markers *SCARA5* and *MFAP5* while also expressing other profibrotic genes *VIM, COL1A2, SPARC*, and *PCOLCE* (*11, 21, 22*) (fig. S1A, B). By the expression these molecular markers, these cells were then designated as fibroblasts. We observed that these cells were in different cellular states that are reminiscent of the various fibroblasts found in murine bleomycin-induced fibrotic and human IPF lungs (*11, 12, 23*) (Fig. 1B, D, E; fig. S1, 3A-C). General fibroblasts (henceforth “fibro” when referring to cell states in our dataset) were enriched for GO terms like ECM organization, mesenchyme differentiation, and TGF-β regulation (fig. S1C). A stress response signature, and expression of *ERO1L, HSPA5*, and *DDIT3*, which are markers of ER stress and the unfolded protein response (*24, 25*), was observed in the cluster fibro ER stress (fig. S1C). Interestingly, we also found a fibroblast population expressing lipid metabolism genes like *PPARG, FABP4*, and *LPL* that we called lipofibroblasts (*26*) (fig. S1C). And finally, we also detected fibroblasts undergoing mitosis and a small population of *IFITM1*^*+*^ fibroblasts enriched for GO terms involving cellular immunity (fig. S1).

### Early-stage IPF ALI-cultures are defined by changes in cellular composition

One of the biggest differences between controls and IPF ALI-cultures can be seen in the shifts of cellular populations (Fig. 2A, B). Basal cells comprise almost a fifth of the entire population in controls; however, their numbers dramatically drop (False discovery rate; FDR adjusted *p*-value < 0.1; table S1) in IPF samples. The opposite observation can be made with the fibro population where their numbers significantly increased in IPF (FDR adjusted *p*-value < 0.1), hinting the progression of fibrosis at the sampling site of the subsegmental bronchus. This expansion of vimentin^+^ fibroblasts was validated in IPF ALI-cultures under low magnification microscopy (fig. S2). Moreover, while the controls showed a regular epithelium, the IPF samples showed a disorganized distribution of cells (fig. S2).

**Figure 2.**
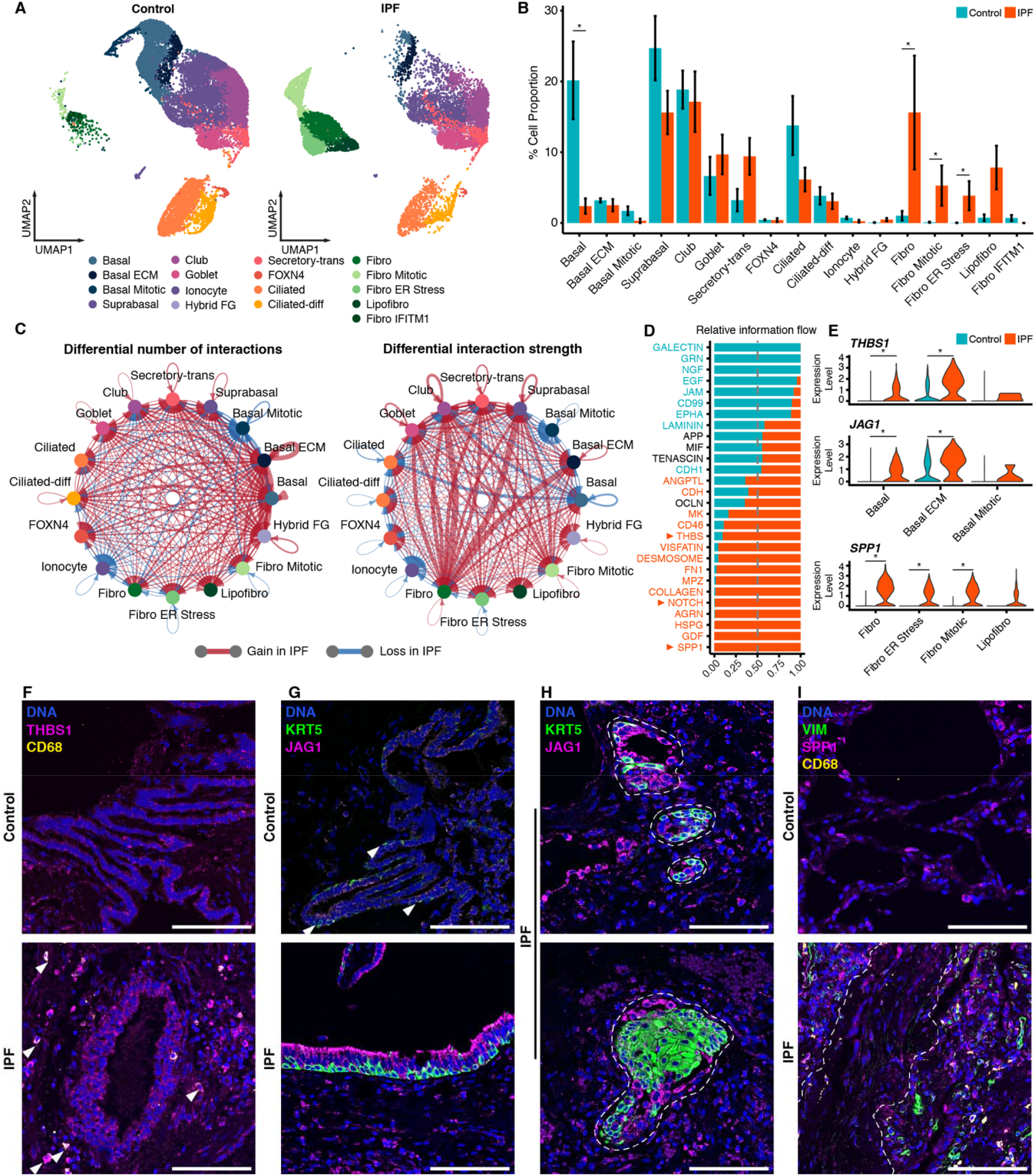
Shifts in early-stage IPF characteristics based on differences in cellular composition, inferred cell-cell interactions, and transcriptional signatures of airway cells. **(A)** UMAP embedding of untreated 17,468 control and 11,457 IPF cells colored by cell type and state annotations. **(B)** Cellular frequencies in percentage of each cell type and state color-coded by disease categorization. Significant differences in cellular populations are indicated by “*” as determined with scCODA with an FDR adjusted *p*-value < 0.1. **(C)** Inferred cell-cell interactions depicting differential interaction counts (left panel) and strengths (right panel) when comparing untreated control and IPF datasets. Edge width indicates the relative interaction count or interaction strength, respectively. Color-coded edges depict a gain (red) or loss (blue) of these metrics in IPF. Fibro *IFITM1* was excluded from the analysis for having too few cells. **(D)** Ranked differences of signaling/interaction pathways. Identified pathways are color-coded according to their significant enrichment to control (cyan) or IPF (orange) conditions as determined by a paired Wilcoxon test. Orange arrowheads highlight pathways of interest. **(E)** Gene expression of highlighted pathway ligands in basal cells (upper panel) and fibroblasts (lower panel) split according to disease conditions. Significant differences are indicated by “*” using the MAST-based differential gene expression test (adjusted *p*-value at < 0.05). **(F-I)** Representative immunofluorescent staining of identified cell-cell interaction mediators and their expression in airway epithelial cells (EPCAM), basal cells (KRT5) and fibroblasts (VIM) in control and IPF lungs. Macrophages are identified by CD68 expression (see fig. S5, 6). **(F)** THBS1 expression by airway epithelial cells and macrophages (white arrowheads). **(G)** JAG1 expression by basal cells (KRT5; white arrowheads) and other airway epithelial cells. **(H)** Basal pod-like structures (outlines) stained with KRT5 and JAG1 in IPF lungs. **(I)** SPP1 expression by fibroblasts and macrophages. Outline highlights region of low SPP1 expressing fibroblasts in IPF lungs. Validation staining: control (n = 2); newly diagnosed IPF (n = 4). Scale bar = 100μm.

### More interactions in early-stage IPF driven by fibroblasts and basal cells

Previous studies of IPF indicated that epithelial-mesenchymal interactions play a key role in the pathogenesis and progression of the disease (*8, 12*); therefore we used CellChat (*27*) to infer how these different cellular compartments may communicate with one another (Fig. 2C, D). Across all cell types, IPF basal cells had the most differential number of interactions, while fibro cells have the highest interaction strength with different epithelial cells like basal ECM, suprabasal, club, and goblet cells (Fig. 2C). To better understand the mechanisms of these interactions, all possible interactions were summarized into specific pathways and their enrichments measured for either controls or IPF (Fig. 2D). ECM-mediated interactions like collagen and fibronectin are enriched in IPF (*11, 12*). We further investigated the SPP1, THBS, and NOTCH pathways as they were shown to be relevant in IPF macrophages (*9, 10*), fibroblasts (*28*), and epithelial cells (*10*). Cells expressing the ligands of these interaction pathways were found to be the different basal cell and fibroblast populations (fig. S4). *THBS1* and *JAG1* were differentially expressed in IPF basal and basal ECM cells, with the latter driving most of the interactions towards other epithelial cells (Fig. 2E; fig. S4). In contrast, *SPP1* was upregulated in IPF fibro, fibro ER stress, and fibro mitotic and interacts mostly with basal cells (Fig. 2E; fig. S4).

Primary lung tissues obtained from newly diagnosed IPF patients were stained for THBS1, JAG1, and SPP1 to validate the enriched IPF interactions (Fig 2F-I; fig. S5). Increased THBS1 expression in basal cells and other airway epithelial cells was observed in IPF lungs (Fig. 2F, fig. S5A). Interestingly, macrophages also highly expressed THBS1, consistent with reports of THBS1 expression in these cells (*29*) (Fig. 2F, fig. S5A). Similarly, JAG1 expression was also increased in IPF airway epithelial cells (Fig. 2G, fig. S5B). Furthermore, across different IPF lungs, basal pod-like structures, of varying sizes that highly expressed KRT5 and JAG1 were observed (Fig. 2H, fig. S6). Finally, SPP1 was mostly expressed by different fibroblasts rather than the sparser macrophages in early-stage IPF lungs. Fibroblasts embedded in collagen expressed SPP1 in lower levels while those surrounding these regions expressed higher SPP1 (Fig. 2I, fig. S5C, D).

### Shared expression of core signatures within epithelial cells and fibroblasts

To better understand differences in the transcriptome, we identified differentially expressed genes per cell type when comparing IPF and controls (fig. S7A; table S2). Within different IPF epithelial cells, multiple proinflammatory chemokines (e.g. *CXCL1, CXCL6, IL8*, and *CCL2*) were upregulated (*30*) (fig. S7A). In addition, we observed increased expression of genes encoding the serum amyloid A (SAA) and S100A family of proteins (e.g. *SAA1, SAA2, S100A8*, and *S100A9*) that are involved in immunomodulation (*31, 32*) (fig. S7A). Apart from inflammation, reparative cytokeratin 6 (*KRT6A* and *KRT6B*) was upregulated by different IPF epithelial cells but mostly by basal cells (*10, 33, 34*) (fig. S7A). In the different fibroblasts, genes involved in motility like the major component of contractile actin *ACTG2*, and an actin-like protein encoded by *ACTBL2* were differentially upregulated in IPF fibroblasts clusters (*11, 28*) (fig. S7A).

We further investigated the expression profile of basal and basal ECM cells with respect to cytokeratin 6 and mediators of TGF-β activation like *THBS1* (fig. S7B). While both basal cell populations tended to show an upregulation of these genes in IPF, basal ECM cells consistently upregulated all the highlighted genes involved in repair processes (*KRT6A, KRT6B*, and *KRT6C*) and TGF-β activation (*THBS1, ITGAV, ITGB5, ITGB6, ITGB8*) (*35*) (fig. S7B).

As we compared the transcriptomes of the different cell types, some genes were consistently upregulated in either IPF epithelial cells or fibroblasts; this prompted us to look at the overlap of the differentially expressed genes (Fig. 3A). In epithelial cells, the comparison was limited to non-transitionary and non-mitotic cell types and excluded rarer cell types like FOXN4 and ionocytes. In this compartment, selected genes were categorized into “core” signatures of immune modulation and *CTNBB1*, encoding β-catenin of the Wnt pathway (Fig. 3B). Here, we again observed immunomodulatory genes like *SAA1, SAA2, S100A8, S100A9*, and *CXCL6* being upregulated by a significant number of epithelial cells that are present in the IPF airway mucosa (Fig. 3B). In newly diagnosed IPF lungs, S100A8/9 was upregulated mainly in airway cell types but was also observed in a few other immune cells (Fig. 3C; fig. S8A). On the other hand, SAA1/2, though expressed in airway epithelial cells of the control lung, were upregulated by many cell types in a more systemic fashion in IPF (Fig. 3D; fig. S8B). Along with this, we also observed highly bronchiolized regions in IPF lungs with increased macrophage infiltrates (Fig. 3E; fig. S8C). Furthermore, the expression of *CTNNB1* was also upregulated by airway epithelial cells (*10, 36*). On the other hand, the core signature of the fibroblasts was mostly composed of genes involved in OXPHOS, and ATP synthesis as revealed by GO term enrichment (Fig. 3A, F, G; OXPHOS gene list was obtained from the enriched GO terms). Once more, *ACTG2* and *ACTBL2* were highly expressed by IPF fibroblasts, along with *AEBP1*, a myofibroblast differentiation enhancer gene (*37*), together defined as “early activation” signature (Fig. 3G).

**Figure 3.**
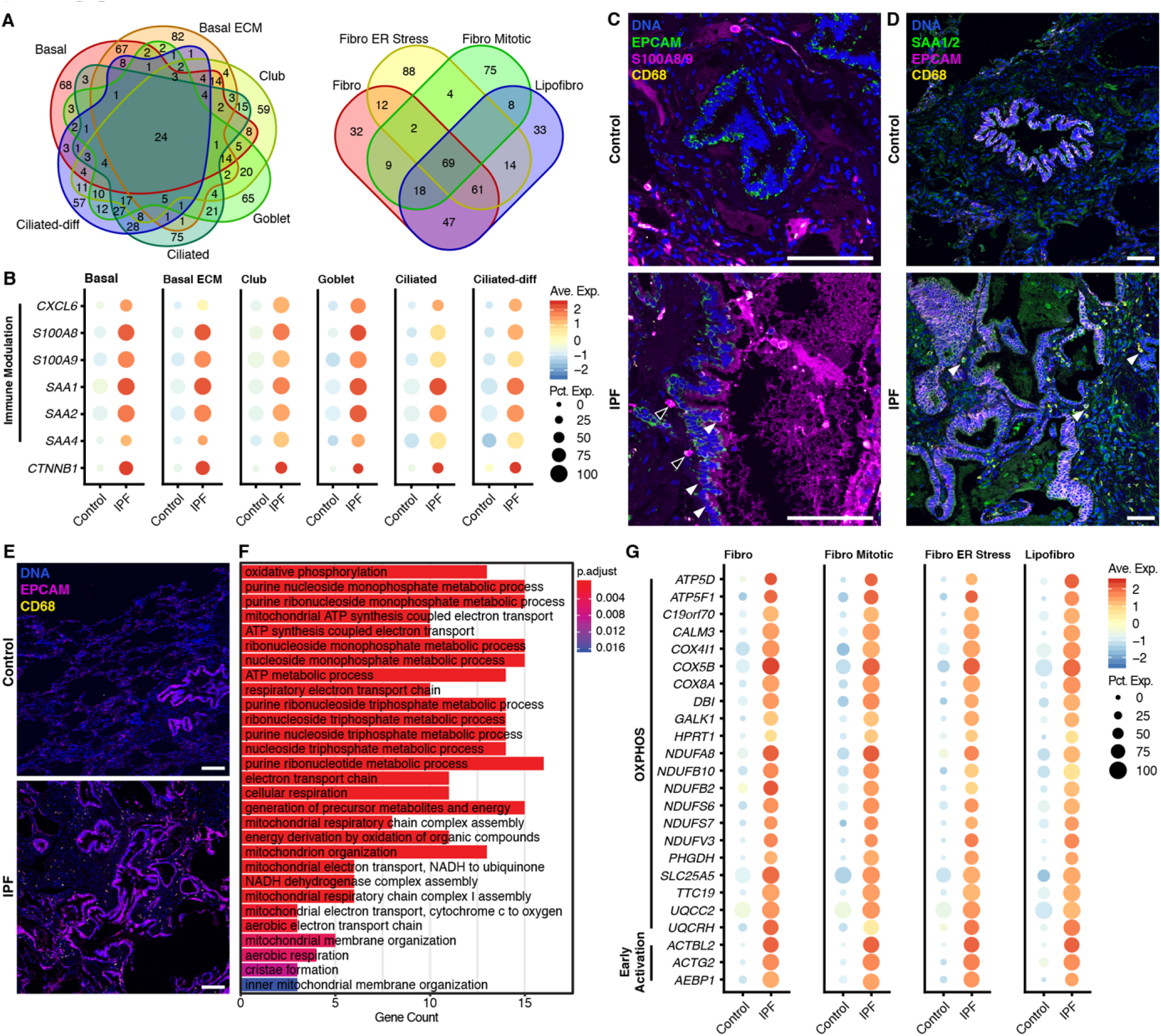
Shared core early-stage IPF signatures in epithelial cells and fibroblasts. **(A)** Overlap analysis of upregulated IPF genes in selected epithelial cells (left panel) and fibroblasts (right panel) to identify “core” IPF genes in each compartment. Epithelial cell selection was limited to cell types that are fully differentiated and are not undergoing a transitionary state. **(B)** Upregulated core genes in highlighted epithelial cell types. **(C-E)** Representative immunofluorescent staining of identified proinflammatory mediators and their expression in airway epithelial cells (EPCAM) in control and IPF lungs. Macrophages are identified by CD68 expression (see fig. S8). **(C)** S100A8/9 expression by airway epithelial cells (white arrowheads) and by other immune cells (black arrowheads) in IPF. **(D)** SAA1/2 expression by airway epithelial cells and other interstitial cells (white arrowheads). **(E)** Increased airway structures in IPF lungs and macrophage presence. **(F)** GO term enrichment of the 69 shared upregulated IPF genes in fibroblasts. Most enriched terms are related to OXPHOS. Significance was determined with a hypergeometric test. **(G)** Upregulated core genes in highlighted fibroblasts. For dot plots: all genes are differentially upregulated in IPF based on MAST-based differential gene expression test (adjusted *p*-value < 0.05). Expression levels are color-coded; the percentage of cells expressing the respective gene is size coded. Validation staining: control (n = 2); newly diagnosed IPF (n = 4). Scale bar = 100μm.

### Treatments are IPF specific and have varying effects on different cells

The current guideline recommends treatment with nintedanib or pirfenidone as these can slow down disease progression (*13*). However, a lung transplant remains the only curative treatment option, if qualified (*1*). Thus, it is imperative to find other drug candidates. To this end, we assessed the effects of saracatinib, an Src kinase inhibitor (*14*–*16*), alongside nintedanib and pirfenidone to investigate their effectiveness in ameliorating the previously described IPF signatures.

The distribution of the different cell types and states remained relatively homogenous after treatment (Fig. 4A; fig. S9A; table S1). To assess the effect of the different treatments, a pseudobulk transcriptome for each cell type from control or IPF conditions for each treatment was generated and used for principal component analysis (PCA; fig. S9B). Cells clustered according to their cell type (ciliated cells along the first component while non-ciliated epithelial cells and fibroblasts along the second component). This unsupervised clustering did not show any separation of clusters driven by disease status or drug treatment. The number of differentially expressed genes was used as a readout to further understand the effect of these three drugs. On the pseudobulk level for each treatment when compared to the untreated control, the IPF samples consistently had more differentially expressed genes (Fig. 4B). This observation remained true even on the single-cell resolution (fig. S9C, table S2). Furthermore, the different fibroblasts enriched in IPF consistently had a stronger response to the three treatments compared to the other cell states (fig. S9C).

**Figure 4.**
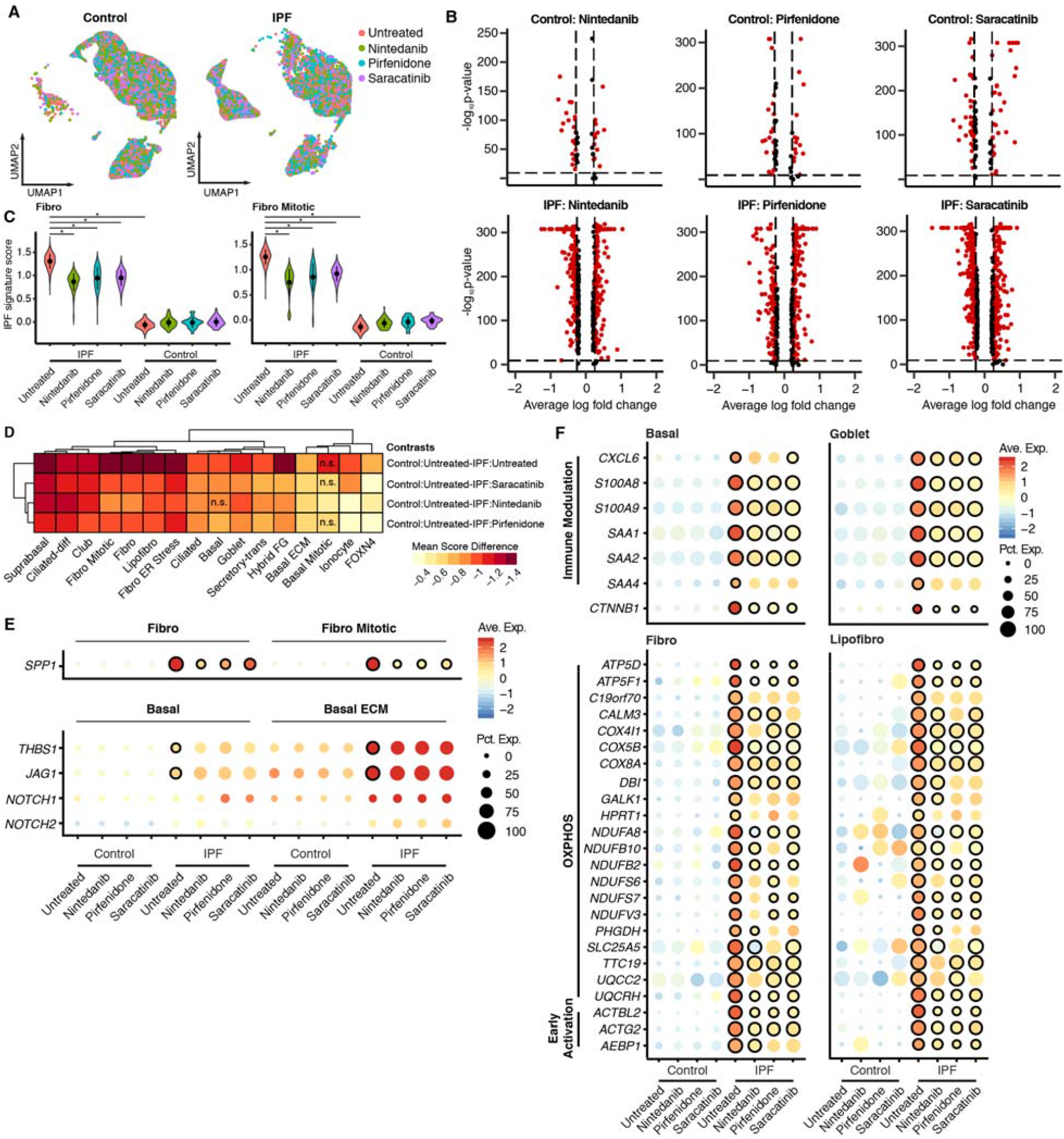
Drug treatments elicit a greater response in IPF cells with varied effects in cell-specific transcription. **(A)** UMAP embedding of ALI-cultures split by control and IPF patients colored by treatment. A subset of 20,000 cells per disease condition. **(B)** Significance (-log_10_ *p*-value) versus magnitude (log fold change) of differentially expressed genes between drug treatments and untreated samples from all cells per treatment group. Dashed lines indicate the log fold change < 0.25 and *p*-value < 0.05 cut-offs. **(C)** Cell-specific IPF signature scores of treatment groups split by disease condition for Fibro and Fibro Mitotic cells (see fig. S7-8). Each dot reflects the mean while the range indicates the standard deviation. Significant differences in IPF signature scores are indicated by “*” using the Dunnet-Tukey-Kramer (DTK) pairwise multiple comparison test (see table S4). **(D)** Heatmap of the mean differences calculated with DTK when comparing the IPF signature scores of untreated control cells across different treated IPF cells. Unless stated otherwise with n.s. = non-significant, observations are significant using the DTK test. **(E)** Upregulated pathway ligands in fibroblasts (upper panel) and basal cells (lower panel) split according to disease and treatment conditions. **(F)** Upregulated core genes in highlighted epithelial and fibroblast cell types/states (see fig. S13). For dot plots: Significance is calculated with a MAST-based differential gene expression test and depicted by a black outline (adjusted *p*-value < 0.05). Significant changes in gene expression were calculated for: untreated normal versus untreated IPF cells, and untreated IPF versus treated IPF cells. Expression levels are color-coded; the percentage of cells expressing the respective gene is size coded.

To investigate the effects of drugs on each cell, we generated a cell-specific IPF signature that is defined as the top 50 genes that are differentially upregulated in IPF (*p*-value < 0.05) with a log fold change of at least 1 (fig. S10). Examples of genes belonging to the cell-specific IPF signature include previously described genes encoding for proinflammatory cytokines (*CXCL1, CXCL6, CXCL5, IL8*, and *CCL20*), immunomodulators (*SAA1, SAA2, S100A8*, and *S100A9*), cytoskeletal components (*KRT6A, KRT6B, ACTG2*, and *ACTBL2*), profibrotic markers (*VIM, FN1, SPP1*, and *MMP7*), and OXPHOS members (*CALM3, NDUFB10*, and *UQCRH*; fig. S10B). Using these cell-specific IPF signatures, a score was calculated for each cell type under the different drug treatments and disease status (see methods; fig. S11; table S3). The differences in the mean of the scores were used as the metric for treatment efficiency, where it is assumed that the treatment should redirect the score towards the control (fig. S11B; table S4). Fibroblasts had the biggest shift in the score after all three treatments compared to the untreated IPF (Fig. 4C, D; fig. S11). We observed a positive shift of the score in the basal ECM cells (Fig. 4D; fig. S11). Comparing the effects of the different drugs on the scores (without looking into specific genes), the shifts were more similar between nintedanib and pirfenidone than with saracatinib as depicted by the hierarchical clustering (Fig. 4D). In general, we observed that different cells have varying degrees of responses to the three drugs.

### Treatments have mixed effects on interaction mediators but significantly decreased core IPF signatures

Next, we assessed if nintedanib, pirfenidone, and saracatinib could affect the pathogenic signatures that we previously described. In terms of cell-cell interactions, it remained that IPF cells were more communicative regardless of the treatment used. Treated IPF cells also showed a higher density of interactions potentially indicating a drug response, where a drug induces the expression of genes involved in cell-cell interactions (fig. S12A). Most probabilities of interaction pathways were minimally changed by the treatment with a few exceptions like NGF, galectin, and GRN (fig. S12B, C). Among SPP1, NOTCH, and THBS pathways, only the SPP1 pathway was significantly affected by the three drugs as shown by the reduced probabilities of the interaction (fig. S12B, C). *SPP1* expression was significantly reduced in all the IPF fibroblasts. However, the interaction partners’ expression levels (*ITGAV, ITGA5, ITGB6*, among others) within basal cells remained largely unaffected by the treatment (Fig. 4E; fig. S13A). On the contrary, *JAG1* and *THBS1* expression were not significantly affected by any of the treatments, and neither was the expression of most of their interaction partners (e.g. *NOTCH1, NOTCH2*, and *NOTCH3* for *JAG1*; *CD47*, and *CD36* for *THBS1*), except for *SDC1* for the THBS pathway (Fig. 4E; fig. S13B). Furthermore, the expression of cytokeratins and TGF-β activation genes in basal and basal ECM cells were also mostly not downregulated by the treatments. The drugs, however, even increased the expression of some genes like *KRT6A* and *ITGB6* in these cells (fig. S13C).

Moving on to the core IPF signatures, it can be observed that most of the genes are significantly affected by the three different drugs, with minimal differences (Fig. 4F; fig. S13D). Within epithelial cells, the expression of immunomodulatory genes *SAA1, SAA2, S100A8*, and *S100A9* was generally downregulated by the three drugs, while *CXCL6* and *SAA4* expression was mostly unaffected. The expression of *CTNNB1* was appreciably downregulated by all three treatments across major cell types (Fig. 4F; fig. S13D). For fibroblasts, the OXPHOS signature was mostly downregulated by three treatments with a few exceptions like *GALK1* and *HPRT1*; similarly, early activation genes *ACTBL2, ACTG2*, and *AEBP1* were all downregulated by the three drugs in all fibroblast cells (Fig. 4F; fig. S13D).

## Discussion

Although previous single-cell transcriptomic studies of human IPF lungs that have identified aberrant cell types and pathogenic transcriptomes, their analyses were limited to understanding end-stage fibrosis by utilizing explant lungs of IPF patients (*8*–*12*). As IPF is a progressive disease, we aimed to study newly diagnosed, early-stage, untreated fibrosis and target the mechanisms that drive its development. In this work, we provide a cellular atlas of the airway mucosa from the subsegmental bronchi that model cellular states during early-stage fibrosis of IPF and their responses to nintedanib, pirfenidone, and saracatinib. With this dataset, we identified characteristics of early-stage fibrosis in IPF airways: (1) dramatic changes in epithelial and fibroblast populations, (2) pathogenic signatures from epithelial cells and fibroblasts, and (3) current pharmacological approaches do not ameliorate all identified signatures of IPF.

### Changes in cellular composition of early-stage IPF ALI-cultures

The biggest difference between controls and IPF ALI-cultures was the shift in cellular populations. This is best exemplified by the reduction of the basal cells and the expansion of the different fibroblasts in IPF samples, which are the only statistically significant changes. However, we do not observe reduced numbers of basal cells in the IPF tissue, and their reduction *in vitro* may potentially reflect the tendency of these cells to differentiate into more differentiated epithelial cells of the mucosa. Finally, we identified an IPF-enriched cell type that is characterized by the shared expression of goblet cell and fibroblast markers, potentially indicating EMT. Previous studies have shown that EMT occurs in IPF where alveolar type 1 (AT1) and AT2 cells can trans-differentiate into fibroblasts (*38*). Here, we identify goblet cells, through hybrid FG cells, as another potential source of fibroblasts by undergoing EMT, though we cannot exclude if this is an artifact of the culture system.

### Epithelial cells initiate local inflammation in early-stage IPF

It has long been suggested that epithelial cells play an important role in IPF as evidenced by genetic alterations and dysregulated gene expression linked to epithelial cells (*5*–*7*). We observed a strong proinflammatory response across the epithelial cells of IPF airways with the expression of SAA, S100, and chemokine genes. The SAA gene family is involved in the acute phase response, which is a non-specific immune response that resolves in 2-3 days; however, they are also involved in chronic inflammation (*31*). Interestingly, SAA1 and SAA2 proteins recruit macrophages and neutrophils and also induce the transformation of macrophages from the proinflammatory M1 state to the tissue reparative M2 state (*31, 39, 40*). The switch to M2-like macrophages is implicated in lung fibrosis (*41*), which is further evidenced by the presence of profibrotic macrophages in IPF lungs (*9*–*11*) and the potential use of monocytes for disease prognostication (*42*). Another set of immunomodulators is the family of S100 proteins that can initiate and amplify inflammation (*32, 43*). Although S100A8 and S100A9 are highly expressed in neutrophils, monocytes, and macrophages, activated epithelial cells in states of inflammation can express them as well (*43*). In this study, we identified for the first time the cells overexpressing these immunomodulatory genes within the airway epithelium of early-stage IPF.

Apart from the shared expression of core immunomodulatory genes, different IPF epithelial cells also upregulated multiple chemokines (e.g. *CXCL1, CXCL6, IL8*, and *CCL2*). These chemokine gradients recruit neutrophils, monocytes, macrophages, and other leukocytes (*30*) and are reflected by the large volume of monocytes and macrophages that are recruited in IPF lungs (*9*–*12*). Evidently, the epithelial landscape of IPF airways plays a significant role in the initiation of fibrosis by promoting local inflammation via proinflammatory cytokine, acute phase response, and inflammation amplifying genes. Together, these proinflammatory mediators may indicate a state of injurious inflammation that normally precedes progressive fibrosis (*41*).

### Early-stage IPF basal cells potentially undergo deleterious repair processes

Basal cells are the stem cell of the airway epithelium and can consequently lead to different respiratory pathologies if gone aberrant. Developmental pathways like Notch and Wnt are essential to the patterning and repair of the respiratory epithelium (*19, 44, 45*). However, prior evidence has shown that they are dysregulated in IPF with increased expression of their mediators like β-catenin, *WNT3A, NOTCH1*, and *JAG1* across epithelial cells including basal cells (*10, 34, 36, 46*). This confirms our observation that both pathways are also dysregulated in early-stage fibrosis basal cells as shown by the enrichment of *NOTCH* interactions and upregulation of *JAG1* and *CTNBB1*.

Keratins are intermediate filaments essential to the maintenance of cellular integrity. Zuo, et al., describe a population of distal airway stem cells (TP63^+^/KRT5^+^) expressing KRT6 that form pod-like structures populated by basal cells (*33*). These basal pods transdifferentiate into AT1 and AT2 cells to regenerate the alveolar epithelium after injury (*33, 47*). Remarkably, we also observe the upregulation of KRT6 genes (*KRT6A, KRT6B*, and *KRT6C*) in our basal and basal ECM cells and observe basal pod-like structures within newly diagnosed IPF lungs highly expressing JAG1.

We hypothesize the local inflammation set by the epithelium leads to the activation of the basal cells (*TP63*^+^/*KRT5*^+^/*KRT6*^+^) to undergo the reparative process. These cells then develop into basal pods within the alveolar space in response to injury of the ongoing fibrosis; however, they fail to undergo alveolarization because of their increased Notch pathway interactions that retains bronchiolar identity (*44*). Ultimately, the basal pods form bronchiolized structures in the alveolar space like honeycomb cysts (*48*). Further controlled experiments need to validate if basal cells lead to the development of bronchiolized honeycomb cysts in IPF, representing failed repair mechanisms, and whether Notch pathway inhibition can rescue this.

### Early-stage IPF basal cells prime the airway epithelium for TGF-β activation

TGF-β is central to the development of IPF because of its roles in proliferation, ECM synthesis, and EMT (*1*). It acts as a chemotactic signal that attracts fibroblasts and promotes their differentiation into activated myofibroblasts (*1*). Sources of TGF-β in IPF include alveolar macrophages, mesenchymal cells, and transformed AT2 cells (*49*). However, as TGF-β is bound to a latency-associated peptide during synthesis, it needs prior activation before eliciting its effects (*35*).

Within the airway epithelium, we observed that the basal ECM cells expressed genes involved in TGF-β activation through various ways. THBS1, the mediator of the THBS pathway, is one of the major activators of TGF-β1 (*50*). A study had shown that IPF airway epithelial cells, but not fibroblasts, expressed *THBS1* (*51*), which is consistent with our observations but now in single-cell resolution. Another way to activate TGF-β is with integrin αv bound to integrin β1/3/5/6/8 (*52*). Our findings that *ITGAV, ITGB5, ITGB6*, and *ITGB8* are upregulated together with *THBS1* in the different basal cells show that these cells are primed to activate TGF-β in early-stage fibrosis.

Aberrant basal cells have been implicated in IPF as shown by their presence in alveolar spaces, honeycomb cysts (*48*), and altered transcriptomes (*11, 34, 48*). Here we highlight the pathogenicity of these cells that initiate repair mechanisms with *KRT6* expression, hijack the developmental pathways Notch and Wnt, and prime the airway epithelium for TGF-β activation by THBS1 and integrins.

### Early-stage IPF fibroblasts are primed for activation and differentiation

The main characteristic of IPF is fibrosis that is brought upon by the activated fibroblasts and myofibroblasts. Studies have shown that there is a wide-variety of fibroblasts in the fibrotic lung (*12, 23*). In our dataset, we observed a similar expansion of diverse fibroblasts. Most of them are undergoing different cellular processes that reflect previously described pathogenic signatures of IPF like cellular proliferation, ECM deposition, ER stress, and lipid metabolism (*1, 53*).

Although *PPARG* expression in lipofibroblasts represses myofibroblast differentiation, evidence has shown that lipofibroblasts can transdifferentiate into myofibroblasts; thus, these cells are seen as a reservoir of potential myofibroblasts (*54*). Similarly, ER stress is tightly linked to the unfolded protein response and they both promote myofibroblast differentiation (*55*). Together, the different cell states indicate that the fibroblast milieu of the subsegmental bronchi in IPF are primed for activation and differentiation.

Within this mesenchymal compartment, we observed a conserved OXPHOS and early activation signature across all enriched IPF fibroblast cells. Although previous studies have shown that reduced OXPHOS activity is a characteristic of fibroblasts (*56*), our data suggest an earlier state of fibroblast activation, where increased OXPHOS could be a compensatory state for increased ECM production. Within the set of early activation genes are *ACTG2, ACTBL2* and *AEBP1*. The first is involved in motility like the major component of contractile actin *ACTA2*, and is upregulated in IPF fibroblasts and myofibroblasts (*11, 28*). While *ACTG2* is a component of the contractile fibers of fibroblasts (*11, 28*), α-smooth actin (α-SMA; encoded by *ACTA2*) is still widely regarded as the bona fide myofibroblast marker (*9, 12*). *ACTA2* is not well expressed by the fibroblasts in this study. ACTBL2 is an actin-like protein involved in growth of certain cancers (*57*). Finally, AEBP1 is a marker of myofibroblast differentiation in IPF, where its expression precedes that of α-SMA and collagens. Taken together, these indicate that the diverse fibroblasts are primed for activation and differentiation.

Osteopontin, encoded by *SPP1*, is a glycoprotein involved in processes like wound healing and inflammation (*58*). With respect to IPF, SPP1 has been shown to be consistently upregulated across many studies (*6, 7*). Recent, scRNA-seq atlases from explant lungs highlight SPP1 expression in profibrotic macrophages, though we observe far more SPP1 expressing fibroblasts than macrophages in newly diagnosed IPF lungs (*9*–*11*). This potentially reflects confounders such as patient diversity and different stages of the disease. In addition, Spp1 was also found to be expressed by profibrotic fibroblasts that are distinct from fibroblasts within fibrotic foci (*59*), which supported by our findings in human IPF lungs. This highlights our observed *SPP1*^+^ fibroblasts from the bronchi as a possible indicator of initiation of fibrosis and progressive remodeling of the IPF airway. In conjunction, these core OXPHOS and early activation signatures along with SPP1 expression highlight the priming of the fibroblasts within the bronchi for activation and differentiation.

### Treatments are specific to IPF cells but do not ameliorate the pathogenic signatures

IPF is currently managed with nintedanib or pirfenidone. Nintedanib is a tyrosine kinase inhibitor that can inhibit several receptors, mainly VEGFR, FGFR, and PDGFR, while pirfenidone has broad-range anti-inflammatory and anti-fibrotic activities like reducing collagen synthesis, TGF-β expression, and fibroblast proliferation (*1, 13*). Though both drugs are able to slow down disease progression, they do not halt disease progression or even reverse fibrosis (*13*). This motivated us to investigate the use of saracatinib as IPF patients have increased activation of Src kinases and Src kinase inhibition was shown to reverse bleomycin-induced fibrosis (*14, 60*). Using these three drugs, we investigated their efficacy in affecting the previously described IPF signatures.

While only minimal changes in cellular densities were observed after treatment, the three drugs induced large transcriptional changes in the IPF cells, suggesting their specificity to the diseased samples and their altered pathways. To measure the effects of the drugs on each cell, we generated a cell-specific IPF signature for each cell type and state and used them to calculate a score for the different treatments. We measured a significant shift of the signatures from the untreated IPF towards the untreated control, almost across all cell types. This also highlights the potential of the use of saracatinib as it has similar shifts in the signatures as nintedanib and pirfenidone. The cells that had the largest shift in the signature are the different fibroblasts that highlights the efficacy of the three different drugs against fibroblasts (*1, 13, 14*). Looking at the pathogenic signatures that we identified, we observed that some of them were significantly affected by treatments while others were not. For example in epithelial cells, we observed that the Notch and THBS1 pathways were not affected whereas the Wnt pathway, represented by β-catenin, was. On the other hand, the SPP1 pathway was significantly affected by the three drugs in the fibroblasts.

Although not all pathways were significantly reduced by the three treatments, what we observed in this study regarding the three treatments reflects complexity of drug effects. Considering that nintedanib, pirfenidone, and saracatinib target different pathways, it may be worthwhile to consider utilizing these drugs in combination with other drugs to maximize effects (*61*).

In summary, we suggest a model that depicts the pathogenic signatures we observe in early-stage IPF into context (Fig. 5). The cascade of events is initiated by different airway epithelial cells that secrete proinflammatory cytokines/chemokines. This subsequently recruits a myriad of leukocytes into the airway mucosa, where monocytes and macrophages further exacerbate the local inflammation. Monocytes and macrophages then secrete latent TGF-β, which will be further processed by basal cells and other airway epithelial cells. All of which are further compounded by the heavy presence of bronchiolized structures in IPF. Once activated, TGF-β elicits its transformative function on the primed fibroblasts within the basal membrane. The activated fibroblasts then proliferate and transdifferentiate into myofibroblasts. Apart from fibrosis, we also hypothesize that IPF basal cells are central to another feature of IPF – the bronchiolized honeycomb cysts. Distal basal cells initiate the repair process by migrating to the damaged alveolar space where they develop basal pods. However, these basal pods fail to transdifferentiate into alveolar cell types due to their increased Notch signaling that retains their bronchiolar identity.

**Figure 5.**
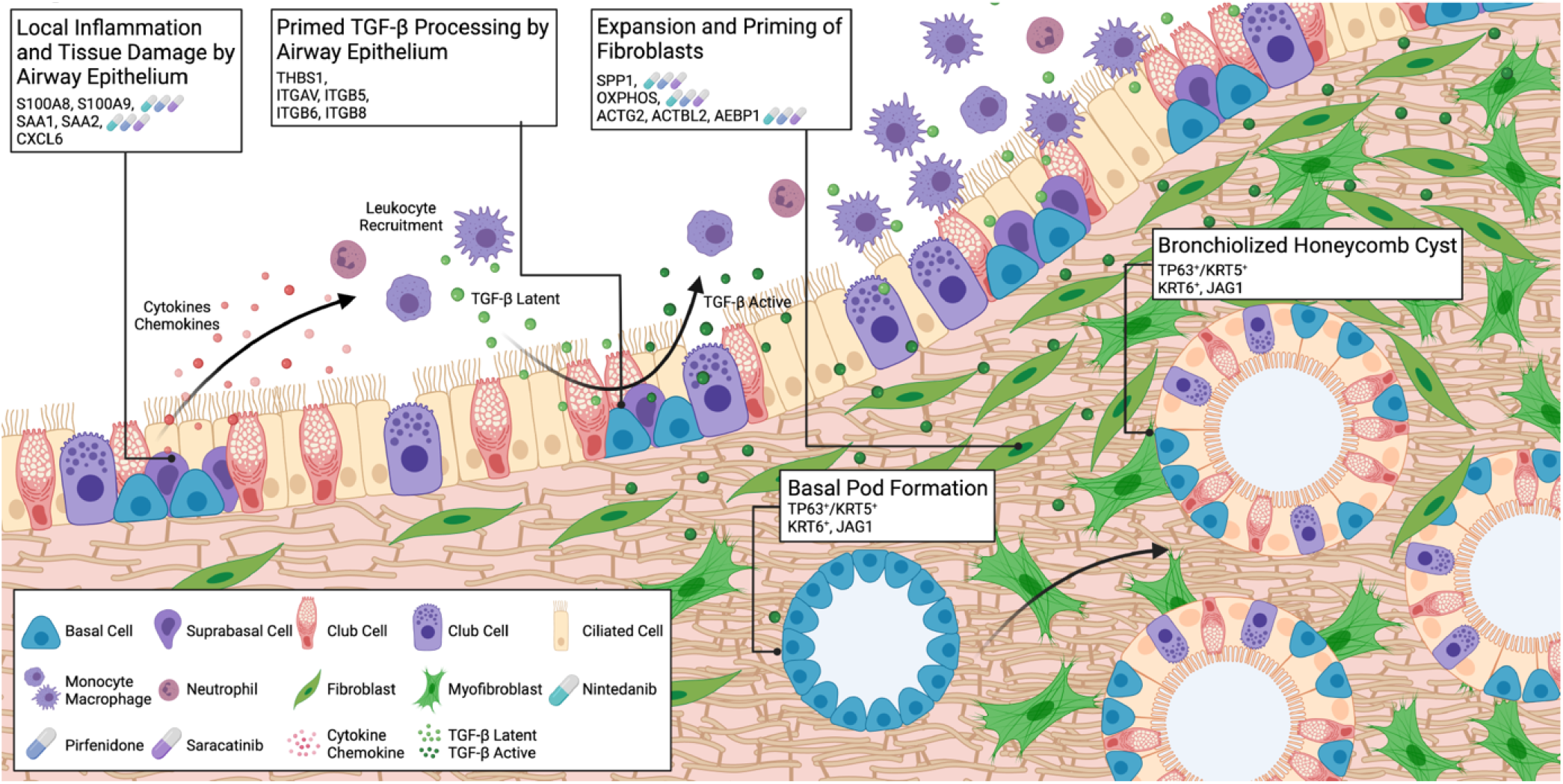
Proposed model of events in early-stage IPF. Airway epithelial cells induce local inflammation and recruit leukocytes. Monocytes and macrophages secrete latent TGF-β that is activated by basal and other airway epithelial cells. Primed fibroblasts are activated and quickly proliferate and transdifferentiate into myofibroblasts that extensively produces ECM proteins. Distal basal cells with high Notch signaling initiate the wound response by migrating and forming basal pods. These fail to transdifferentiate and retain their bronchiolized identity and form the bronchiolized honeycomb cysts in distal ends of the IPF lung. The legend of the icons is depicted in a box. Mediators marked with a drug icon indicate that they are downregulated by the specific treatment (nintedanib – cyan; pirfenidone – blue; saracatinib -purple).

It must however be noted that our study also comes with its limitations. The first is that our observations are limited to our sample size and future studies would benefit from having larger cohorts. Second, we cannot rule out any changes in the transcriptome that is brought about by the culture model. Thus, any potential findings derived from our dataset must always be validated in primary tissue. In addition, the observed presence of fibroblasts within the ALI-culture system was perplexing but can be explained by the lack of selection methods used after acquiring cells with bronchial microsampling. This allowed us to obtain a heterogenous pool of cells that included HBECs and airway fibroblasts that were then used to generate the ALI-cultures. Other studies have also shown the ability of the ALI medium to support the growth of fibroblasts (*62, 63*). Lastly, the use of a 24-hour treatment period allowed us to study short-term effects of these drugs but further studies should include extended timepoints to evaluate any long-term effects of the treatments.

Here, we provide a rich dataset that investigates the early-stage transformation of proinflammatory epithelial cells, pathogenic basal cells, and primed fibroblasts from the medial airways of IPF and find that nintedanib, pirfenidone, and saracatinib only affects some of the identified signatures in our study but miss to inhibit primed basal cell populations as potential drivers of fibrosis development. This study provides more insight into the disease mechanisms of IPF and might serve as a key resource for the IPF community to further investigate the effects of potential pharmacological inhibition strategies.

## Materials and Methods

### Patient recruitment and ethics approval

All patients provided informed consent to take part in the study following the Declaration of Helsinki, and the Department of Health and Human Services Belmont Report. The Medical Faculty of Heidelberg ethics committee approved the use of the biomaterials for this study (S-270/2001 and S-538/2012).

### Air-liquid interface (ALI) cultures of HBECs

Primary human bronchial epithelial cells (HBECs) were obtained from the ELF of patients by minimally invasive bronchoscopic microsampling from subsegmental airways as previously described (*64*). Patient characteristics are available in table S5.

Primary HBECs were cultured following previously published methods (*18*). Briefly, cells were grown to confluence before transferring to a 12-well plate with ThinCert™ cell culture inserts with 0.4μm pores (Greiner Bio-One, 665641). Cells were airlifted after 2-3 days by removing the medium from the apical chamber and providing PneumaCult-ALI media (StemCell, 05021) in the basal chamber only. At day 28, HBECs were treated with 0.1µM nintedanib (Selleck Chemicals, S1010), 1mM pirfenidone (Selleck Chemicals, S2907), and 1µM saracatinib (Selleck Chemicals, S1006) for 24 hours. These concentrations were based on plasma levels of each compound from patients, where nintedanib is around 0.1µM and pirfenidone is about 100µM(*65*–*68*). However, pirfenidone is well tolerated *in vitro* and so a higher concentration was used for our purposes(*65*). For saracatinib, the lowest concentration that reduced the phosphorylation of Src family kinases at tyrosine 416 was selected (data not shown). For all drug concentrations used in this study, cell viability and cell death were assessed by MTT and LDH assays (data not shown).

### Dissociation and cryopreservation

To generate single-cell suspensions, the cells were washed three times with 1X PBS (Merck, D8537) to remove excess mucus. Normal samples were dissociated with 0.25% Trypsin-EDTA (ThermoFisher, 25200056) for 10 minutes followed by Dispase (Corning, 354235) for another 10 minutes at 37°C. Diseased samples were treated with 0.25% Trypsin-EDTA for 15 minutes at 37°C. Proteases were deactivated with culture medium. Cells were spun down at 300*g* for 5 minutes, resuspended in 1X PBS, and passed through a 20μm cell strainer (pluriSelect, 43-50020-03). Samples were then stored in cryopreservation medium (10% DMSO, 10% FBS, and 80% PneumaCult-ALI media) and gradually frozen down to -80°C before being transported on dry-ice.

### Generating scRNA-seq libraries

Cryopreserved cells were thawed at 37°C, spun down at 300*g* for 5 minutes, resuspended in 1X PBS with 0.05% Bovine Serum Albumin (BSA; Merck, 05479), and passed through a 35μm filter (Corning, 352235). Single-cell suspensions were loaded following the protocol of the Chromium Single Cell 3’ Library Kit v2 (10× Genomics; PN-120237, PN-120236, PN-120262) to generate cell and gel bead emulsions. The succeeding steps of reverse transcription, cDNA amplification, and sequencing library generation were all performed according to the manufacturer’s instructions. Libraries were sequenced with NextSeq500 (Illumina; high-output mode, paired-end 26x49 bp) or HiSeq4000 (Illumina; high-output mode, paired-end 26×74 bp). Previously published ALI-cultured controls were included in this study (EGAS00001004419: SAMEA6848752, SAMEA6848753, SAMEA6848754, SAMEA6848755) (*18*).

### Pre-processing and analysis of scRNA-seq data

Raw sequencing data were processed using Cell Ranger software version 2.1.1 (10x Genomics). Transcripts were aligned with the 10x reference human genome hg19 1.2.0. Pre-processing to remove low-quality cells was performed with Seurat v3.1.5 based on the following criteria: (i) >200 and <3000-5000 genes, (ii) <15% mitochondrial reads (*69, 70*) (fig. S14). Canonical correlation analysis (CCA) was used for integration and batch correction. PCA and UMAP were calculated and finally, clustering was performed. Contaminating ambient RNA reads were removed with DecontX which is included in the celda package v1.6.1 (*71*).

Cell type assignment was performed as previously described (*18*) and additional cell types were determined. Comparison of cell-type identity with previously published IPF scRNA-seq cell atlases was done using the label transfer function implemented in Seurat as TransferData(). Potential cell-cell interactions between the different cell types were identified using CellChat v0.5.5 (*27*). Differential interactions between patient groups were calculated using mergeCellChat() and compareInteractions() functions iteratively per comparison. CellChat generates an intercellular communication network that is a weighted directed graph composed of significant interactions between cell groups. “Interaction strength” is defined as the communication probability of each computed network while “information flow” is the overall communication probability, where it is the summation of the probability among all pairs of cell groups in the inferred network (*27*). Monocle3 was used for trajectory graph learning and pseudoLJtime measurement through reversed graph embedding (*72*). Cell-specific IPF signatures were generated by selecting the top 50 differentially expressed genes in untreated controls and IPF samples with thresholds of a log fold change of >1 and *p*-values of < 0.05. These were then used to calculate the cell-specific IPF score using the function AddModuleScore() from Seurat. Gene ontology (GO) term enrichment was performed by using clusterProfiler v3.12.0 enrichGO() function (*73*). Differentially expressed genes used as input had adjusted *p*-values < 0.05. Euler plots were generated with the venn v1.10 package in R.

### Immunofluorescence staining

ALI-culture inserts were stained as previously described (*18*) (table S6 for antibody list). In short, filters were fixed in 4% paraformaldehyde (PFA; Merck, P6148) and permeabilized with 0.1% Triton X-100 (Merck, 93443). Primary antibodies were incubated overnight at 4°C. Samples were then applied with secondary antibodies for 40 minutes at 37°C. Nuclei were stained with Hoechst 33342 (Merck, B2261). Micrographs were taken with a Leica SP5 confocal microscope.

Lung biopsies from IPF patients were obtained by video-assisted thoracoscopic surgery (VATS) and embedded in paraffin. The IPF tissue sections were then checked for the presence of different proteins (table S5-6 for patient and antibody lists). Tissue sections were deparaffinized and immersed in 1X sodium citrate buffer (Merck, C9999) kept at 95°C for 10 minutes. Subsequent steps follow the instructions of VectaFluor™ Excel Amplified Kits (Vector Laboratories; DK-1488, DK-2594). To remove autofluorescence, tissue sections were sequentially treated with Vector TrueVIEW™ Kit (Vector Laboratories, SP-8400), and Lipofuscin Autofluorescence Quencher (PromoCell, PK-CA707-23007) following their respective described protocols. The sections were then stained with 16μM Hoechst 33258 (ThermoFischer, H3569) for 5 minutes and washed with 1X PBS before mounting with VECTASHIELD® HardSet™ Antifade Mounting Medium (Vector Laboratories, H-1400-10). Stained slides were then visualized with a Leica SP8 confocal microscope. All images were processed and assembled with FIJI (*74*).

### Statistical Analysis

Statistically significant differences in cellular proportions were calculated with scCODA (FDR adjusted *p*-value < 0.1) (*75*). Differential gene expression testing was calculated using FindMarkers() with a MAST-based differential expression test from Seurat and significance was set at *p*-value < 0.05. When multiple tests were performed, *p*-values were adjusted using the Benjamini-Hochberg method and considered significant when adjusted *p*-value < 0.05. CellChat identifies enriched pathways to a specific condition by performing a paired Wilcoxon test internally within the rankNet() function. ClusterProfiler implements a hypergeometric test to determine significance and thresholds were set for Benjamini-Hochberg adjusted *p*-values at < 0.05. Significant differences in IPF signatures were computed with a Dunnet-Tukey-Kramer pairwise multiple comparison test using the DTK package v3.5 (*76*). Additional visualization packages were also used in this study: pheatmap v1.0.12 (*77*), EnhancedVolcano v1.2.0 (*78*), ComplexHeatmap v.2.0.0 (*79*).

## Data Availability

Raw sequencing data are uploaded in the European Genome-phenome Archive under the reference number EGAXXXXXX. Accessing the data requires a Data Transfer Agreement.

## Acknowledgments

We thank the patients who kindly agreed to donate tissue samples to make this study possible. We also thank David Ibberson (CellNetworks Deep Sequencing Core Facility, Heidelberg University) for sequencing services. Last, we would like to thank Sören Lukassen and Luca Tosti for the helpful discussions regarding single-cell analysis. Overview and summative figures were created with BioRender.com.

## Author contributions

Project Conceptualization and Design: NCK, CC

Drug Treatment Design: CV, AWB

Primary Tissue Provision: NCK, CV, AWB, ECX

Experimentation: RLC, CV, MAS, KJ, ECX

Data Analysis: RLC Supervision: CC, NCK

Writing—original draft: RLC, CC, NCK

Read, Review, and Editing: RLC, NCK, CC, CV, MAS, KJ, ECX, AWB, MK, RE

