## Supplemental Figures for "Profibrotic priming of airway cell types and drug responses in early-stage idiopathic pulmonary fibrosis"

#### Supplementary Figure 1

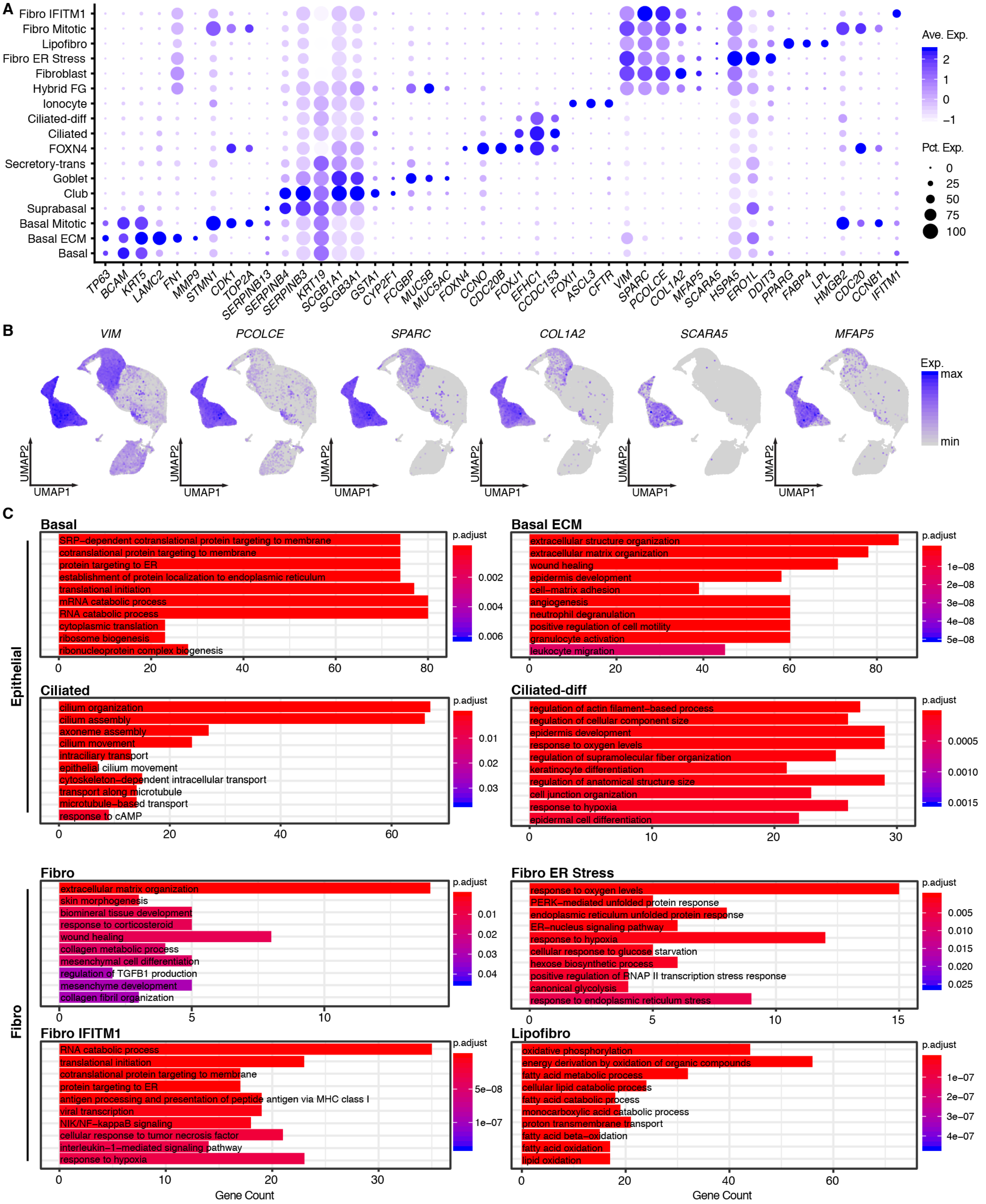


**Figure S1. Marker genes and enriched GO terms used to define the different cell types. (A)** Marker genes used to annotate the different cells. Expression levels are color-coded, and the percentage of cells expressing the respective genes are size coded. **(B)** UMAPs showing fibroblast markers. **(C)** GO term enrichment for subclusters of major cell types. Significance was determined with a hypergeometric test (adjusted *p*-value < 0.05).

#### Supplementary Figure 2

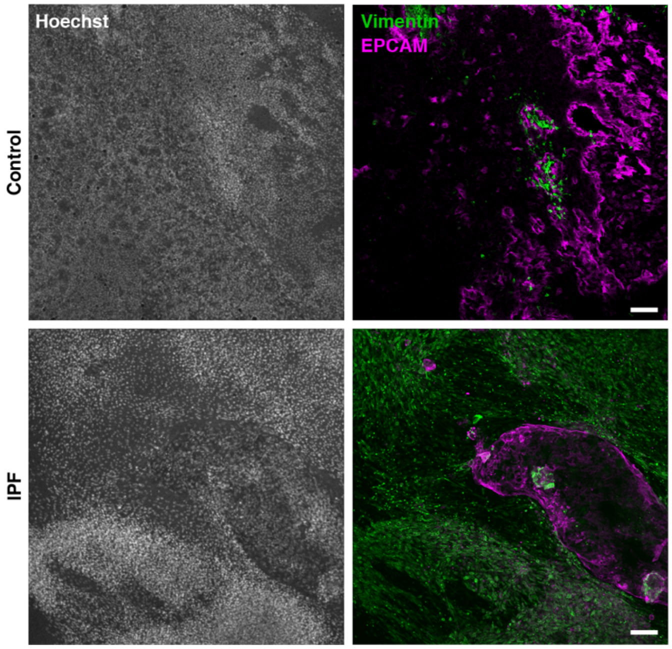


**Figure S2. Overviews of ALI-cultured cells from controls and early-stage IPF patients.** Low magnification overview of ALI-cultures highlighting epithelial and fibroblast composition. Validation cultures: control (n = 2); newly diagnosed IPF (n = 2). Scale bar = 100µm.

#### Supplementary Figure 3

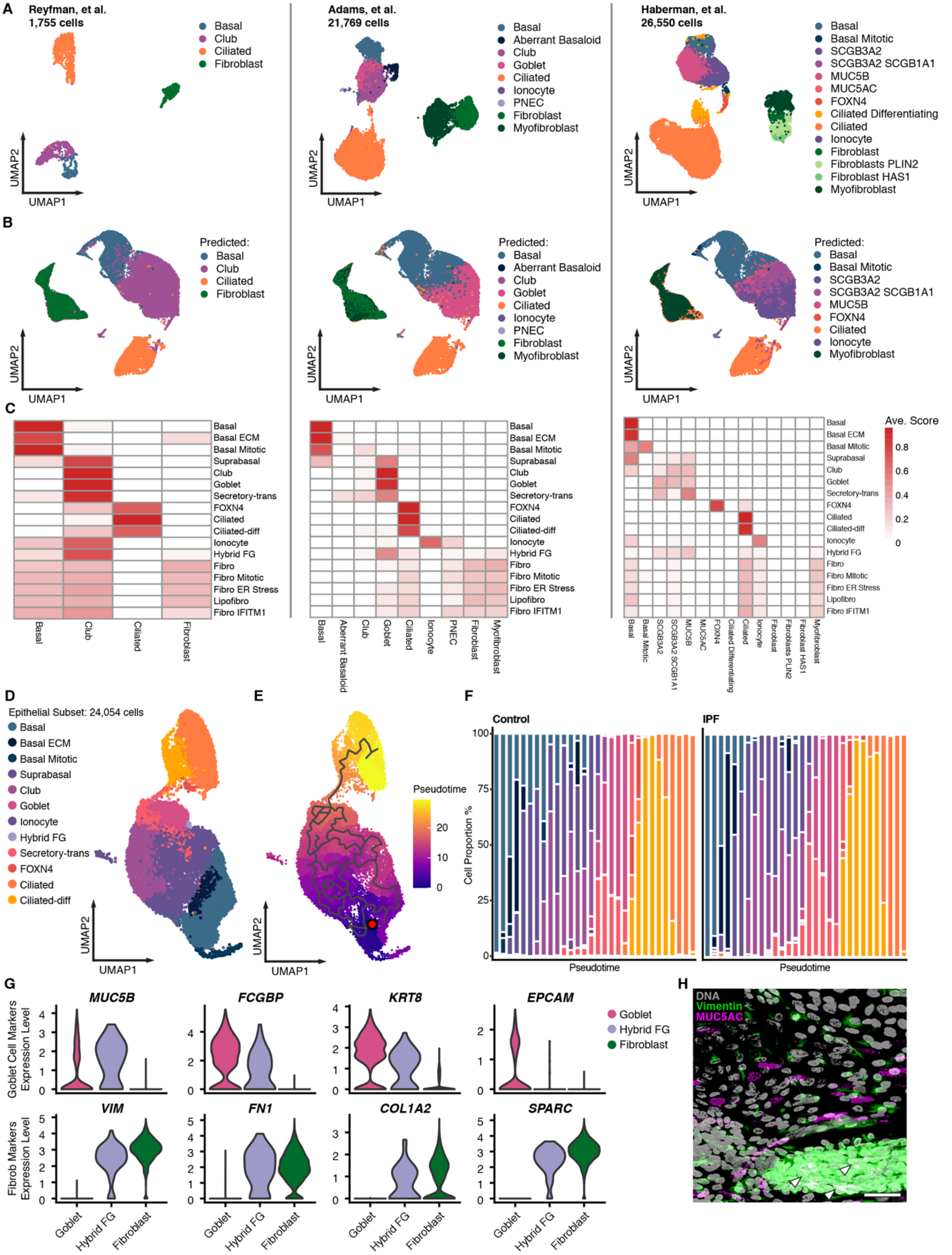


**Figure S3. Reference comparisons, pseudotemporal dynamics and hybrid cells.** Publicly available scRNA-seq IPF atlases generated from control and IPF lungs were subsetted for airway epithelial cells and fibroblast cells. **(A)** UMAP embedding of airway epithelial cells and fibroblasts of each single-cell atlas. Cell type identity is color-coded for each dataset. The Reyfman dataset was reconstructed from count matrices to the final annotation. The other datasets are subsets from their respective publicly available single-cell objects. **(B)** Predicted cell type identity after performing label transfer using other IPF cell atlases as reference. **(C)** Heatmap of the mean prediction score per cell type. **(D)** UMAP of the epithelial subset from untreated control and IPF samples. **(E)** Pseudotime trajectory projected on a UMAP of the subset of all epithelial cells from untreated control and IPF samples. Calculated pseudotime values are color coded. The red dot indicates the start point of the trajectory. **(F)** Cell type composition along a binned pseudotime axis (left to right). **(G)** Expression of goblet (top panel) and fibroblast (bottom panel) markers in goblet, hybrid FG, and fibroblast cells. **(H)** Representative immunofluorescence staining of fibroblast clusters in IPF ALI-cultures. White arrowheads highlight MUC5AC protein located within the fibroblast clusters. Scale bar = 50µm. Validation cultures: control (n = 2); newly diagnosed IPF (n = 2).

### Supplementary Figure 4

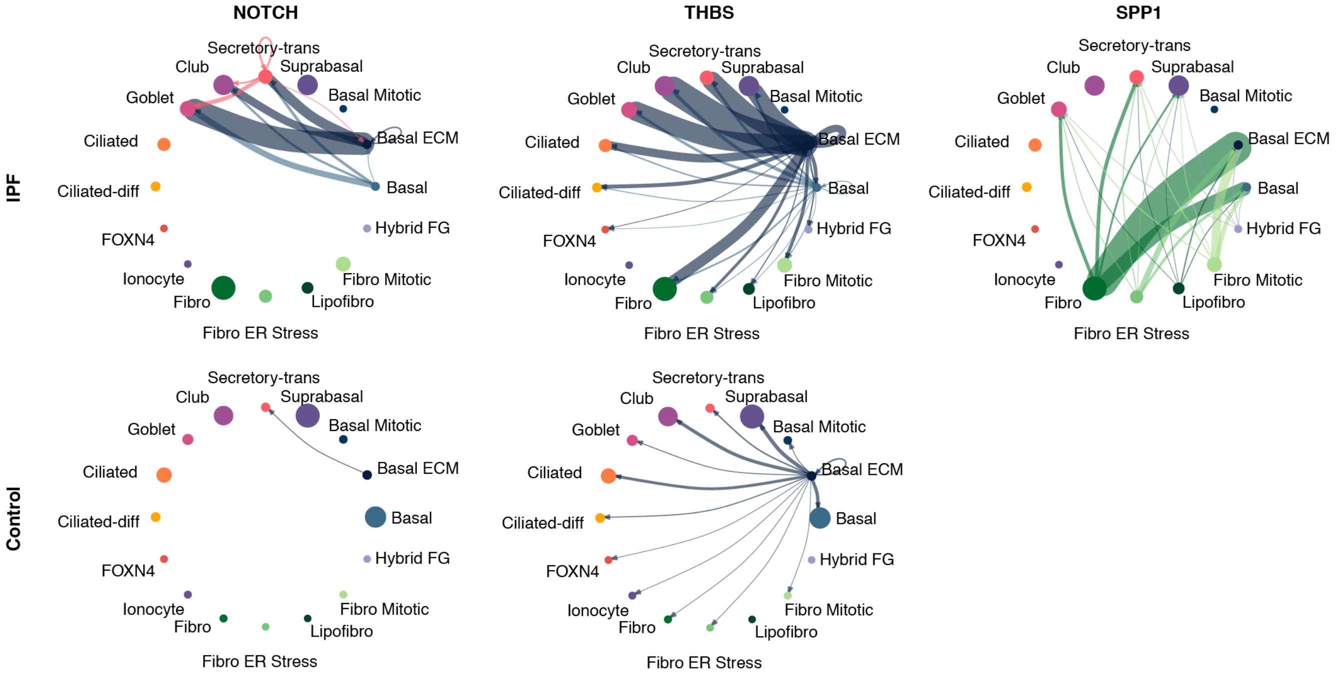


**Figure S4. Inferred cell-cell interactions in IPF cells.** Circle plots depicting cells involved in highlighted signaling/interaction pathways enriched in IPF. Edge colors indicate the sender of the interaction while edge weights are proportional to interaction strength. Edge weights are comparable only within each pathway. Dots are proportional to the number of cells. The SPP1 interaction pathway is not detected in the normal condition.

#### Supplementary Figure 5

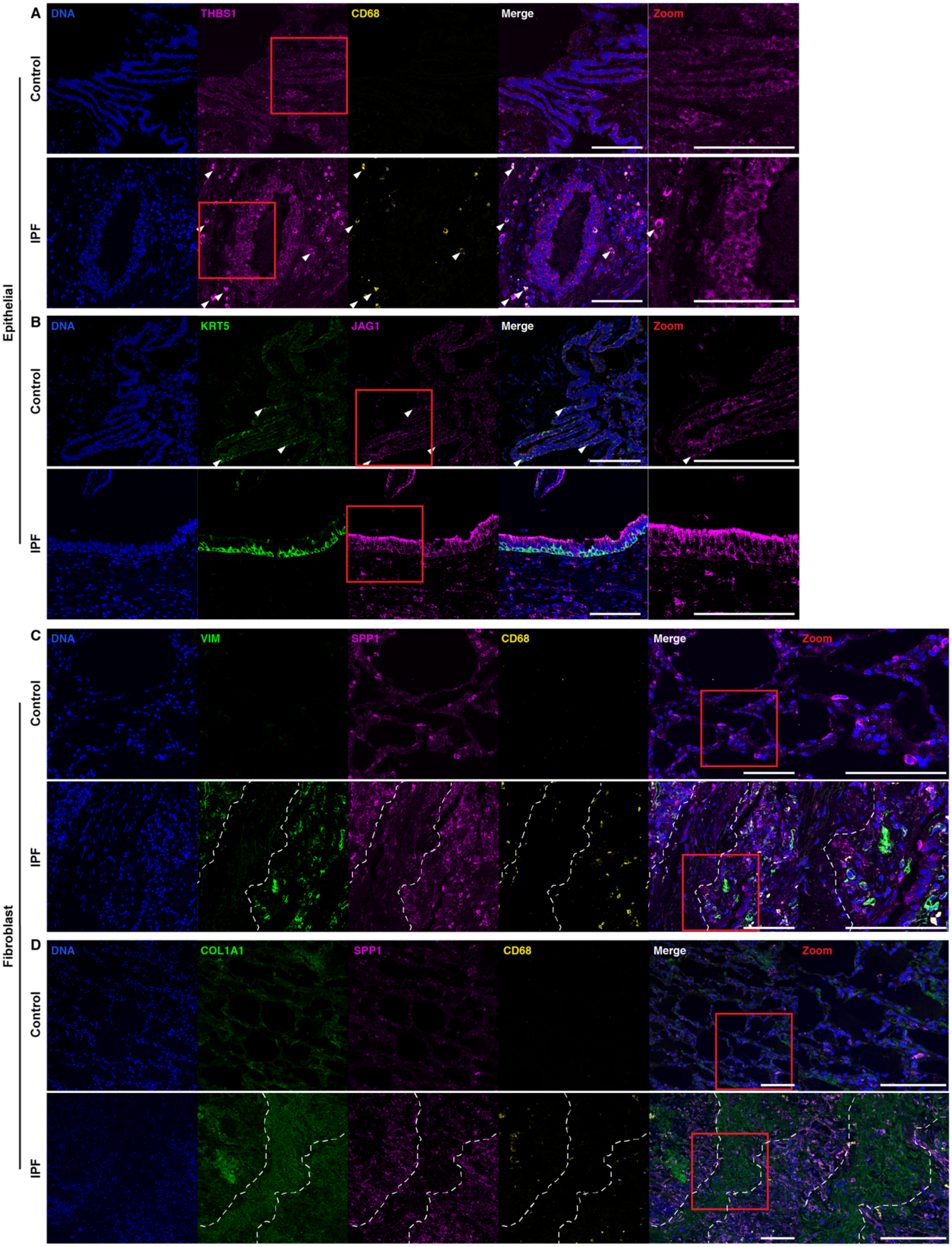


**Figure S5. Primed TGF-β processing, increased JAG1 expression, and activated fibroblasts in early-stage IPF.** Immunofluorescent staining of identified pathogenic signatures within airway epithelial cells (EPCAM) and fibroblasts (VIM) in control and IPF lungs. Macrophages are identified by CD68 expression. **(A)** THBS1 expression by airway epithelial cells and macrophages (white arrowheads). **(B)** JAG1 expression by basal cells (KRT5; white arrowheads) and other airway epithelial cells. **(C)** SPP1 expression by fibroblasts and macrophages. Outline highlights region of low SPP1 expressing fibroblasts. **(D)** SPP1 expression in fibroblasts varies and is associated with collagen deposition (outline). Validation staining: control (n = 2); newly diagnosed IPF (n = 4). Scale bar = 100µm.

#### Supplementary Figure 6

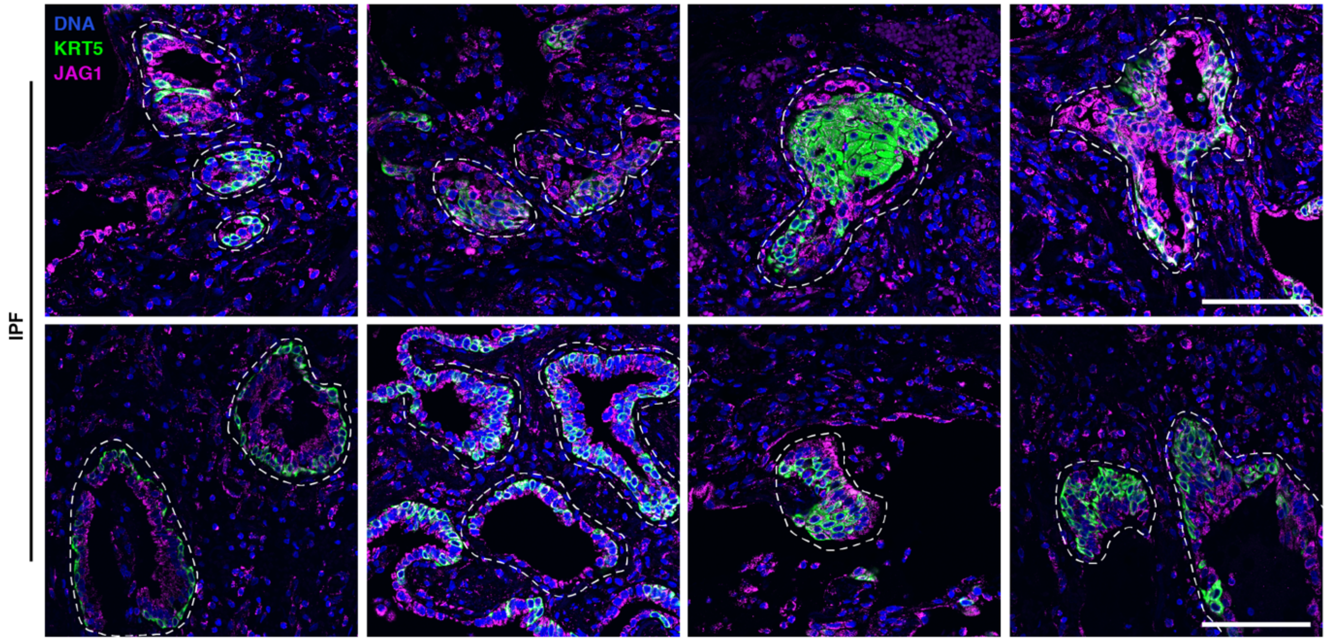


**Figure S6. Basal pod-like structures in early-stage IPF.** Immunofluorescent staining of varying sizes of basal pod-like structures (outlines) stained with KRT5 and JAG1 in IPF lungs. Validation staining: newly diagnosed IPF (n = 4). Scale bar = 100μm.

### Supplementary Figure 7

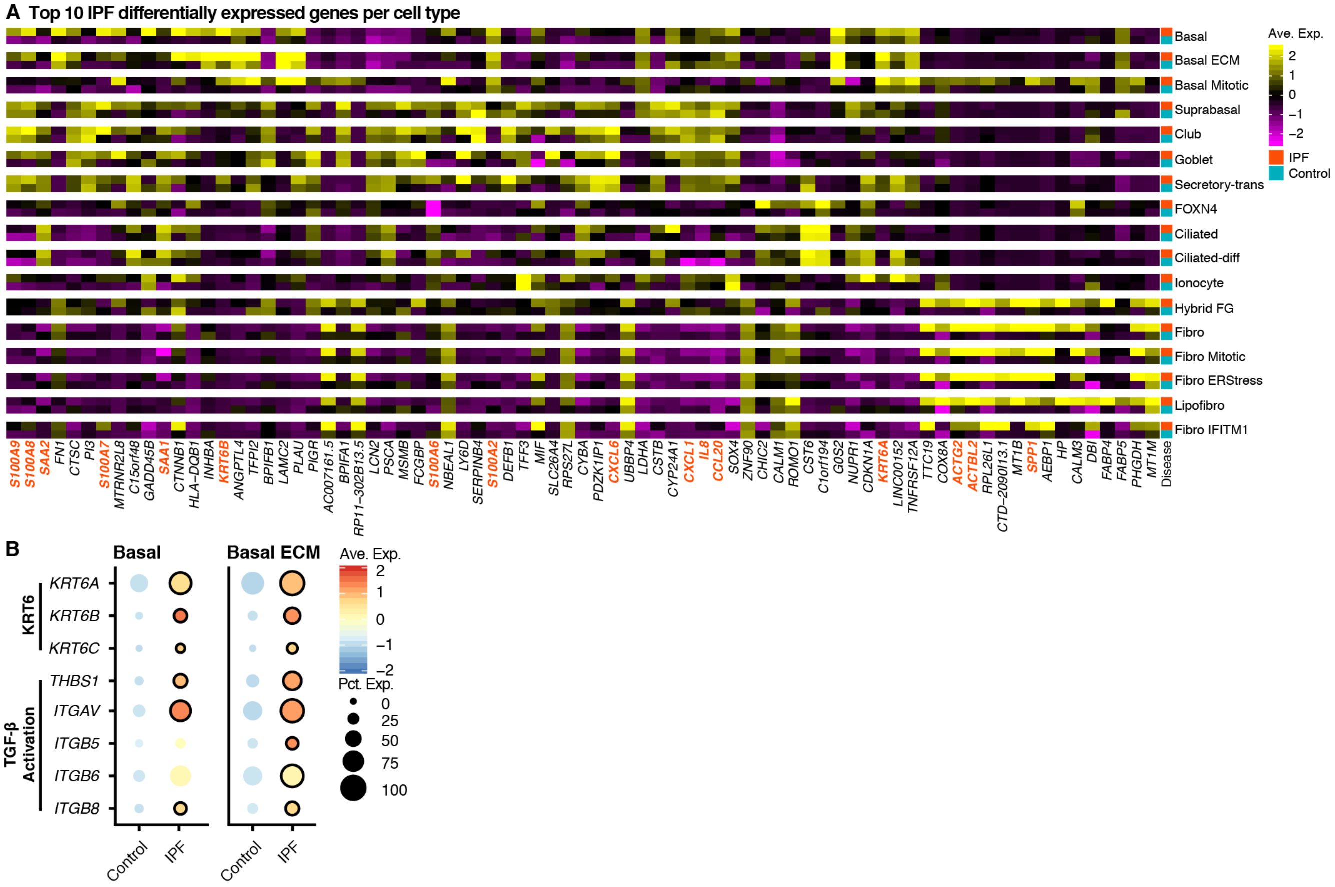


**Figure S7. Differentially expressed genes across cell types. (A)** Top 10 upregulated IPF genes of each cell type/state versus normal samples. Bars are color-coded according to disease conditions. Differential gene expression testing was performed with a MAST-based test (adjusted $p$-value at $< 0.05$; see Supplementary Table 4). Genes in orange are discussed in text. **(B)** Expression of cytokeratin 6 and TGF-β activation genes in basal and basal ECM cells. Significance between untreated control versus untreated IPF cells is depicted by a black outline. Expression levels are color-coded; the percentage of cells expressing the respective gene is size coded.

#### Supplementary Figure 8

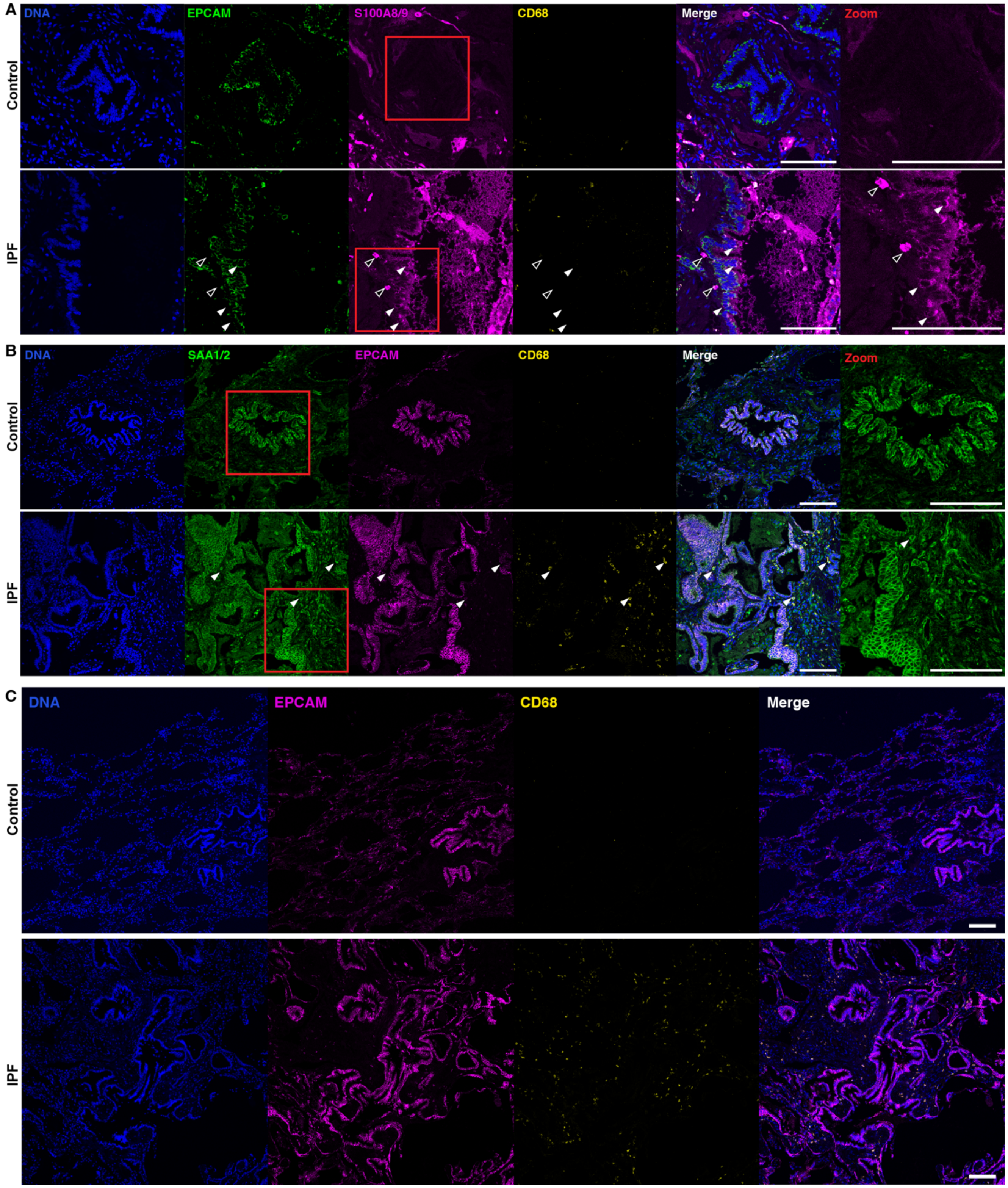


**Figure S8. Proinflammatory epithelial cells and bronchiolization in early-stage IPF.** Representative immunofluorescent staining of identified proinflammatory mediators and their expression in airway epithelial cells (EPCAM) in control and newly diagnosed IPF lungs. Macrophages are identified by CD68 expression. Regions with increased magnification are in a red box. **(A)** S100A8/9 expression by airway epithelial cells (white arrowheads) and by other immune cells (black arrowheads) in IPF. **(B)** SAA1/2 expression by airway epithelial cells and other interstitial cells (white arrowheads) of the IPF lung. **(C)** Increased airway structures in IPF lungs and macrophage presence. Validation staining: control (n = 2); newly diagnosed IPF (n = 4). Scale bar = 100µm.

#### Supplementary Figure 9

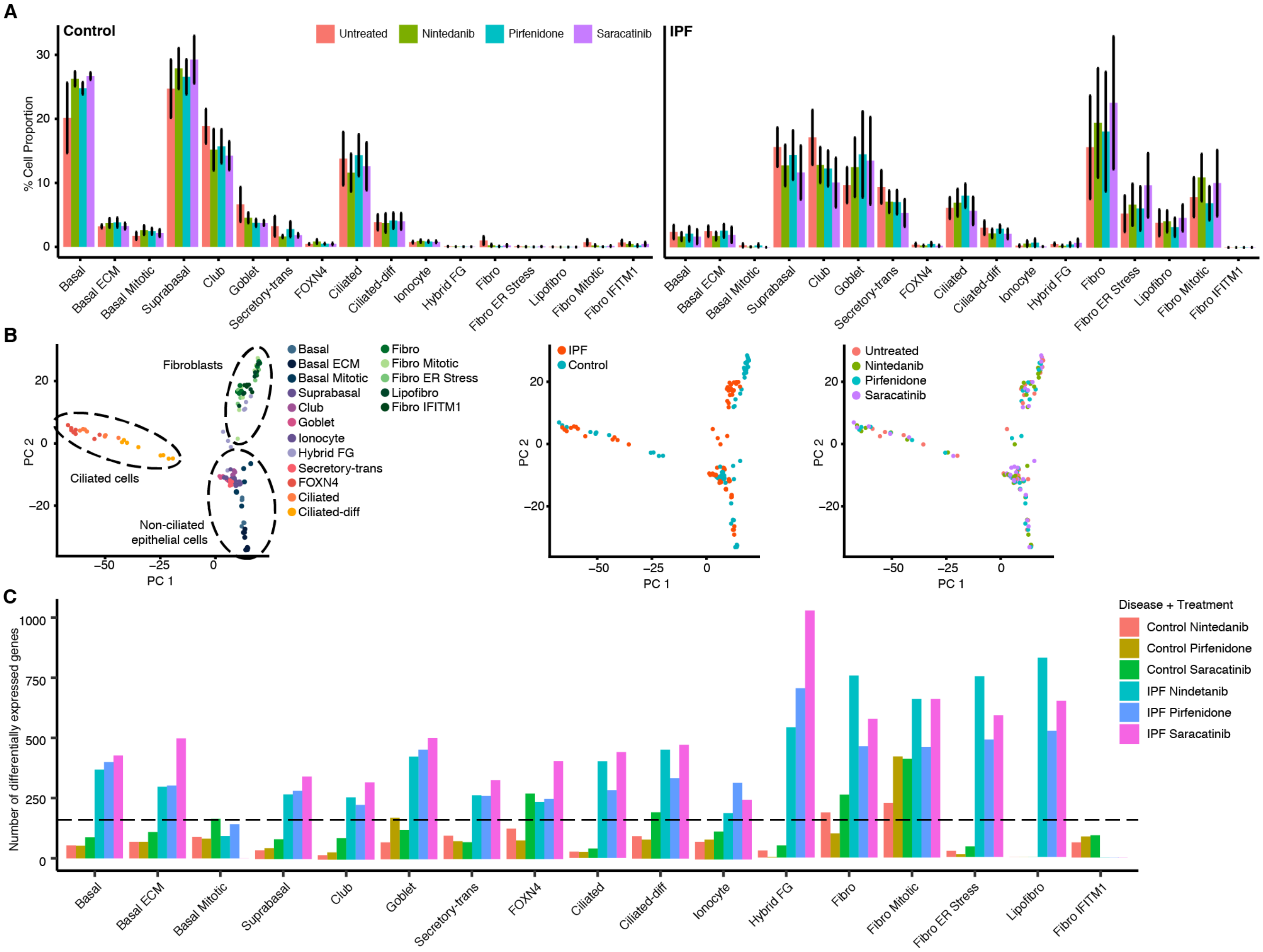


**Figure S9. Treatments do not change cellular populations but affect IPF cells transcriptionally. (A)** Cellular frequencies in percentage of each cell type/state for control (left panel) and IPF (right panel) samples color-coded by treatments. No significant differences in cellular populations as determined with scCODA. **(B)** PCA of pseudobulk transcriptomes of each cell type per disease condition per treatment color-coded according to cell type (left panel), disease (middle panel), and treatment (right panel). Dashed circles highlight the clustering of non-ciliated epithelial cells, ciliated cells, and fibroblasts cell types with each other. **(C)** Number of differentially expressed genes for each cell type when comparing untreated with treated samples in either control or IPF samples. Differential gene expression was performed with a MAST-based test. The dashed line at 200 emphasizes higher numbers of differentially expressed genes in IPF cells.

#### Supplementary Figure 10

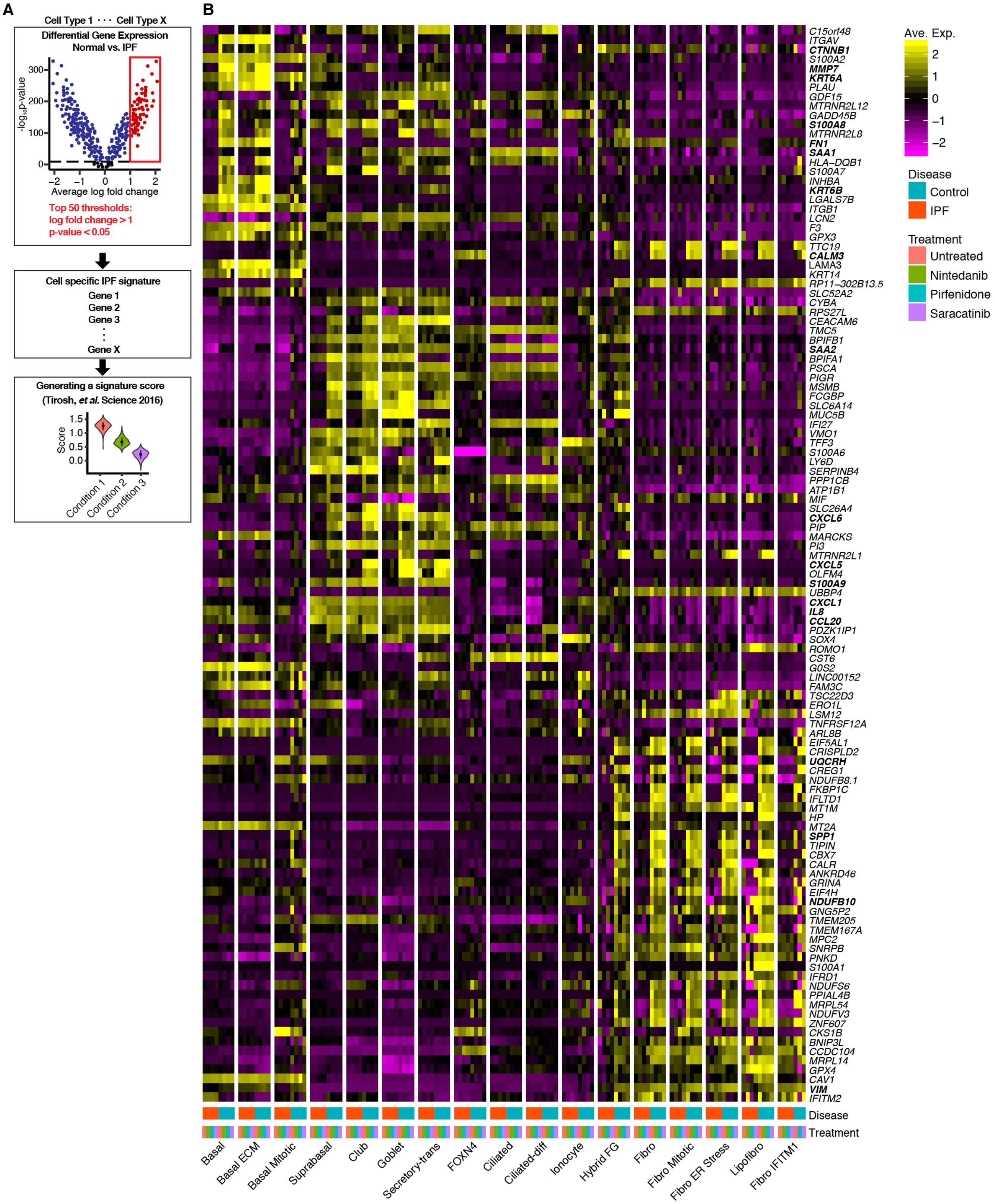


**Figure S10. Generating a cell-specific IPF signature. (A)** A schematic to depict the selection of cell-specific IPF signatures and computation of the score. **(B)** Heatmap of cell-specific IPF signatures (Supplementary Table 5). Colored bars indicate disease and treatment categories.

### Supplementary Figure 11

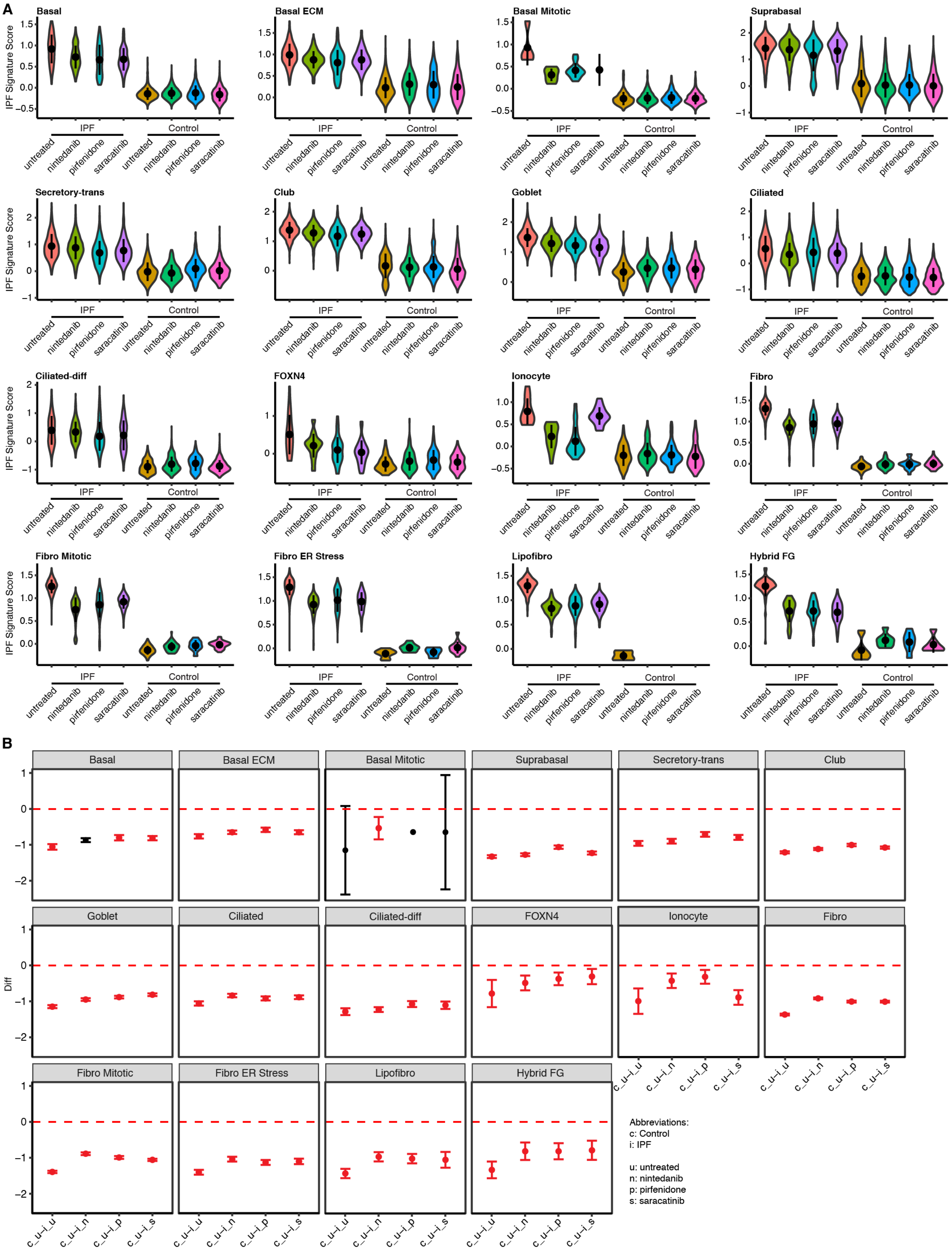


**Figure S11. Cell-specific IPF signature scores highlight significant differences upon treatment. (A)** Calculated cell-specific IPF signature scores split according to disease and treatment conditions. Each dot reflects the mean while the range indicates the standard deviation. **(B)** Dunnet-Tukey-Kramer (DTK) pairwise multiple comparison test confidence interval plots for selected comparisons (Supplementary Table 6). The dot indicates the mean while the range reflects the upper and lower confidence intervals. Significance is color-coded where significance differences are depicted in red, and non-significant differences are in black.

### Supplementary Figure 12

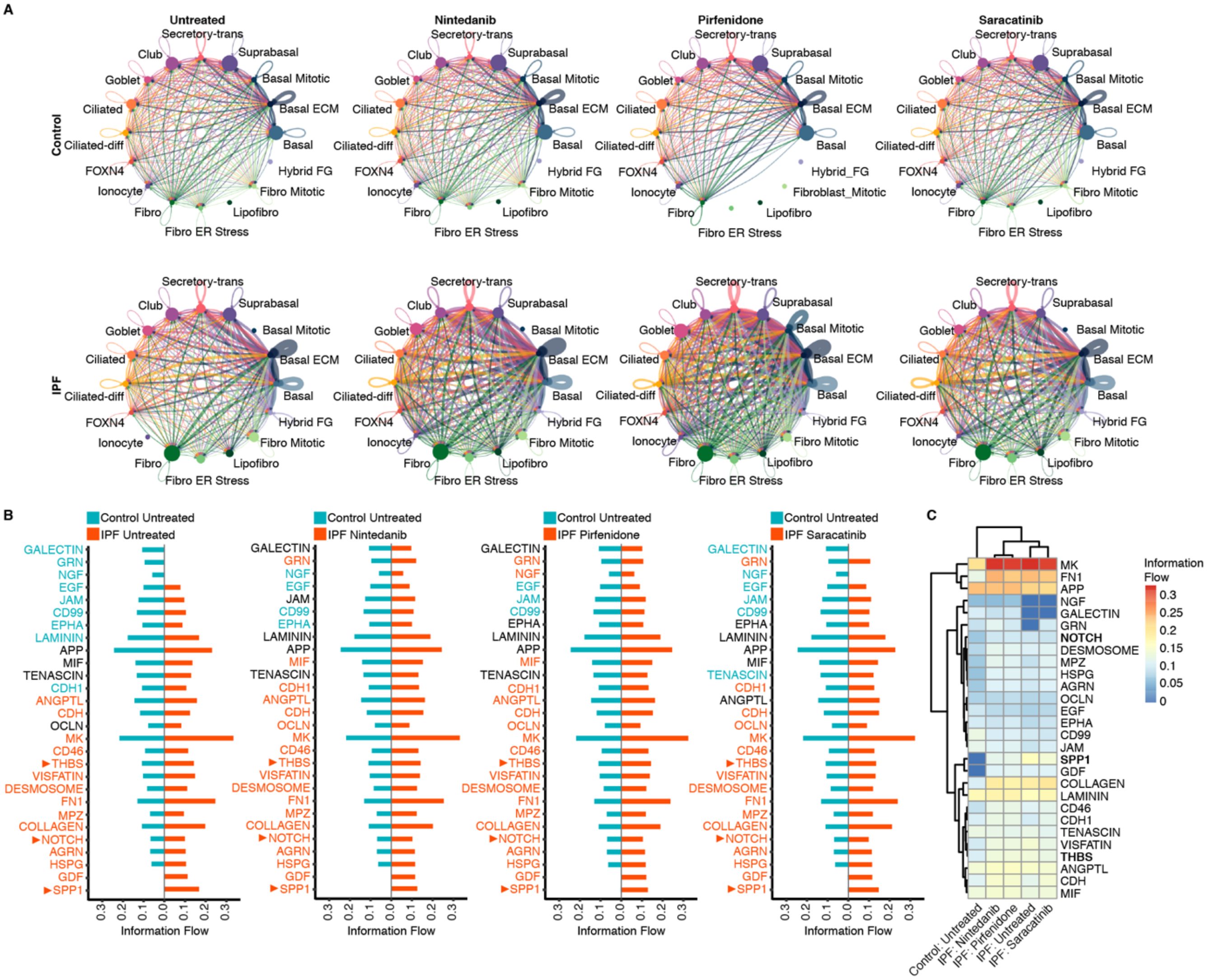


**Figure S12. Cell-cell interactions after treatment. (A)** Aggregated cell-cell communication networks for each disease condition and treatment. Edge colors indicate the sender while edge weights are proportional to total interaction strength. Dots are proportional to the number of cells. **(B)** Differences of signaling/interaction pathways. Identified pathways are color-coded according to their significant enrichment to a specific condition as determined by a paired Wilcoxon test. Pathways are listed following the ranking of untreated normal and IPF comparison. **(C)** Heatmap of the calculated information flow for each pathway in subfigure B with hierarchical clustering.

#### Supplementary Figure 13

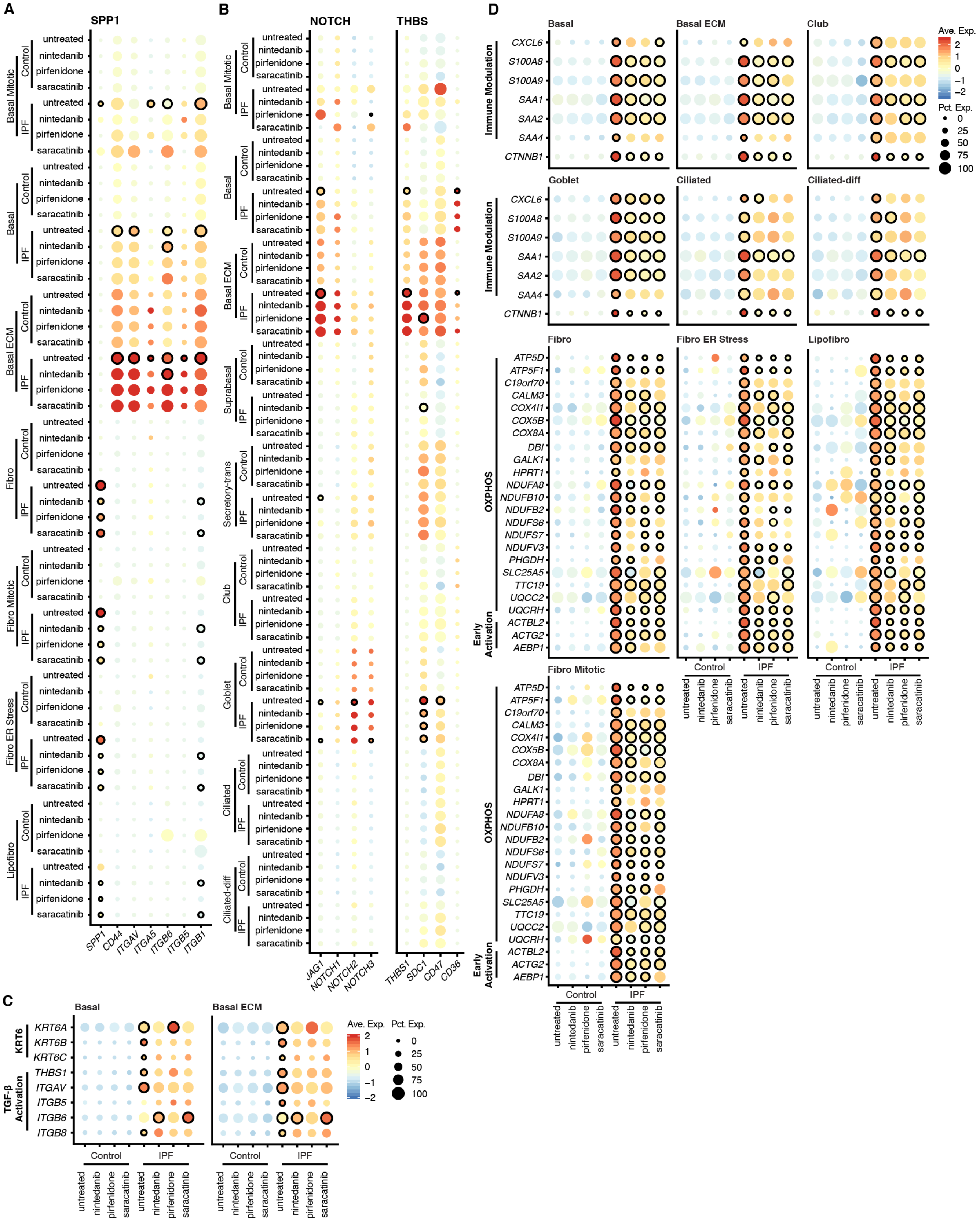

**Figure S13. Gene expression profiles of enriched interaction pathways and core IPF genes. (A-B)** Genes involved in SPP1, NOTCH, and THBS signaling/interaction pathways as identified with CellChat. Cells selected for plotting were based on Supplementary Figure 4A. **(C)** Expression of cytokeratin 6 and TGF-β activation genes in basal and basal ECM cells. (**D)** Upregulated immune-modulatory and Wnt genes by highlighted epithelial cell types and OXPHOS and actin-related genes in fibroblast cell types/states. For dot plots: significance is calculated for treated IPF conditions with the untreated IPF sample.

Significance is calculated with a MAST-based differential gene expression test and depicted by a black outline ($p$-value < 0.05). Significant changes in gene expression were calculated for: untreated normal versus untreated IPF cells, and untreated IPF versus treated IPF cells. Expression levels are color-coded; the percentage of cells expressing the respective gene is size coded.

### Supplementary Figure 14

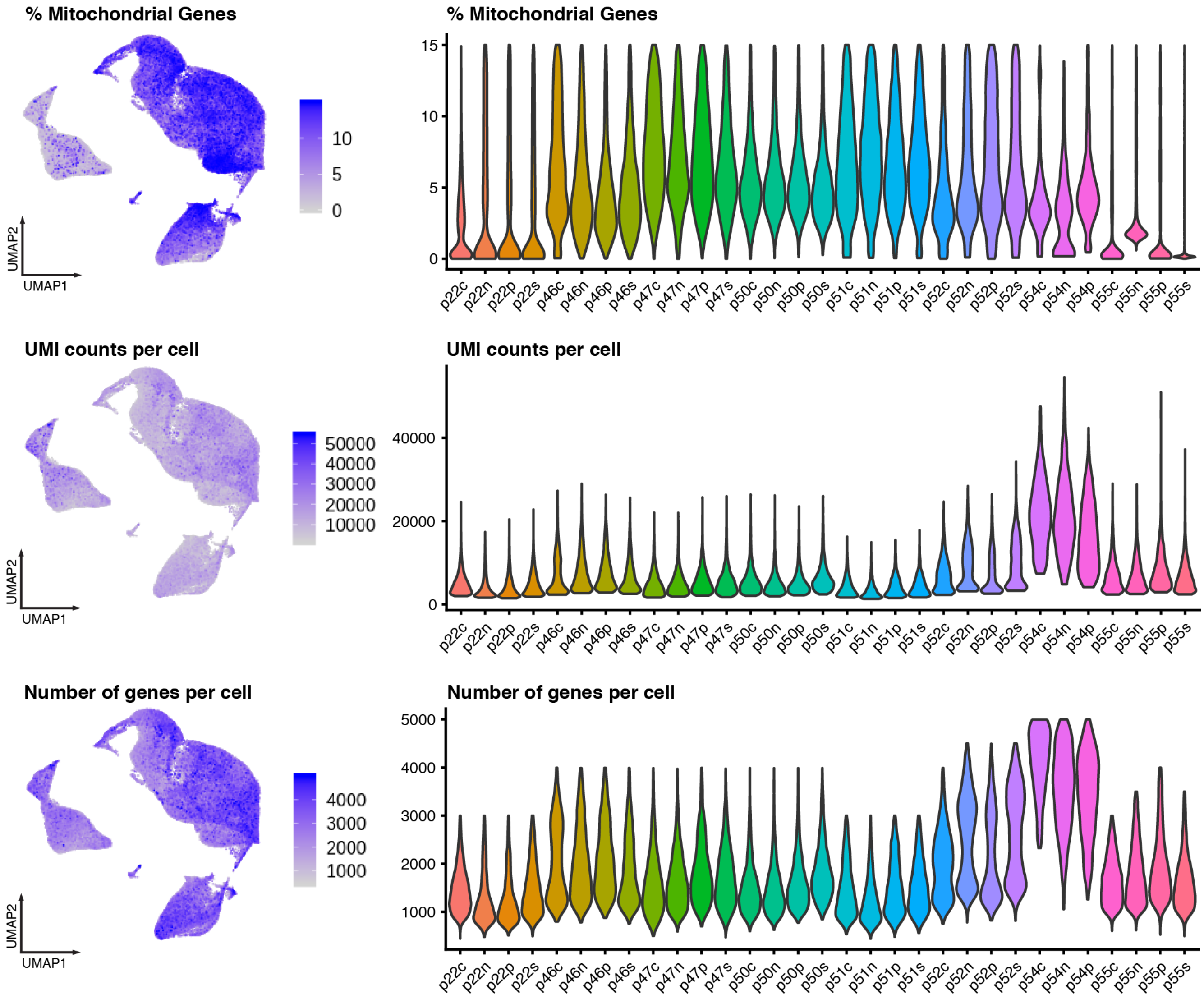


**Figure S14. Quality control metrics.** UMAP and violin plots depicting **(A)** percent mitochondrial genes, **(B)** UMI counts, and **(C)** total genes detected per cell across all samples. Abbreviations: c – control, n – nintedanib, p – pirfenidone, s – saracatinib.

**Table S1. Statistics on cellular population shifts by disease and treatment using scCODA.** (separate file).

**Table S2. Differentially expressed genes per cell type/state by disease and treatment.** Differential gene testing was performed with a MAST-based test**.** For multiple test comparisons, the *p*-values were adjusted with the Benjamini-Hochberg method. Differences in expression are considered significant when the *p*-value or adjusted *p*-value (for multiple comparisons) are $< 0.05$. (separate file).

**Table S3. Genes comprising the cell-specific IPF signatures.** The signature is composed of the top 50 differentially expressed genes that passed the log fold change of $> 1$ and *p*-values of $< 0.05$ using a MAST-based test. (separate file).

**Table S4. DTK pairwise multiple comparison test for changes in IPF signature scores with respect to the treatments.** (separate file). PY – pack years.

**Table S5. Overview of relevant patient characteristics.** DLCO – diffusing capacity of lung for carbon monoxide; $FEV_1$ – forced expiration at the first second; FVC – forced vital capacity. (separate file).

| Antibody | Host | Dilution | Amplified? | Sample | Brand and Catalog Number |
|---|---|---|---|---|---|
| SPP1 | Mouse | 1:100 | Yes | VATS | Invitrogen, 14-9096-82 |
| S100A8/9 | Mouse | 1:100 | Yes | VATS | Santa Cruz Biotech, sc-48352 |
| SAA1/2 | Rabbit | 1:100 | Yes | VATS | Thermo Scientific, PA5-112852 |
| JAG1 | Mouse | 1:100 | Yes | VATS | Santa Cruz Biotech, sc-390177 |
| THBS1 | Mouse | 1:100 | Yes | VATS | Santa Cruz Biotech, sc-59887 |
| KRT5 | Rabbit | 1:100 | No | VATS | Biozol, ATA-HPA059479 |
| EPCAM | Mouse | 1:100 | No | VATS | ThermoFischer, MA5-12436 |
| EPCAM | Rabbit | 1:100 | No | VATS | Thermo Scientific, PA5-19832 |
| Vimentin | Rabbit | 1:100 | No | VATS | Abcam, ab92547 |
| COL1A1 | Rabbit | 1:100 | No | VATS | Cell Signaling, 72026T |
| CD68 | Rat | 1:100 | No | VATS | Abcam, ab53444 |
| ACE2 | Rabbit | 1:100 | No | VATS | Abcam, ab15348 |
| TMPRSS2 | Mouse | 1:100 | Yes | VATS | Sigma, HPA035787-100UL |
| Goat α-Rabbit AlexaFluor®488 | Goat | 1:500 | - | VATS | Life Technologies, A11034 |
| Goat α-Mouse AlexaFluor®568 | Goat | 1:500 | - | VATS | Life Technologies, A11004 |
| Goat α-Rat AlexaFluor®647 | Goat | 1:500 | - | VATS | Life Technologies, A21247 |
| FABP4 | Goat | 1:50 | - | ALI | R&D Systems, AF3150 |
| ERO1L | Mouse | 1:50 | - | ALI | Santa Cruz Biotech, sc-365526 |
| SCGB1A1 | Rabbit | 1:300 | - | ALI | Biozol, BVD-RD181022220-01 |
| B-tubulin IV | Mouse | 1:50 | - | ALI | Sigma, T7941 |
| MUC5AC | Mouse | 1:300 | - | ALI | Abcam, ab3649 |
| KRT5 | Rabbit | 1:50 | - | ALI | Biozol, ATA-HPA059479 |
| Vimentin | Rabbit | 1:100 | - | ALI | Abcam, ab92547 |
| Donkey α-Rabbit AlexaFluor®488 | Donkey | 1:300 | - | ALI | Dianova, 711-545-152 |
| Donkey α-Mouse AlexaFluor®549 | Donkey | 1:300 | - | ALI | Dianova, 715-545-150 |
| Donkey α-Goat AlexaFluor®549 | Donkey | 1:300 | - | ALI | Dianova, 705-585-147 |

**Table S6. Primary and secondary antibodies used for immunofluorescence.** Antibodies whose signals were amplified with the VectaFluor™ Excel Amplified Kits are indicated. VATS – Video assisted thoracic surgery; ALI – air-liquid interface culture.
